# Nutations in plant shoots: endogenous and exogenous factors in the presence of mechanical deformations

**DOI:** 10.1101/2020.07.06.188987

**Authors:** Daniele Agostinelli, Antonio DeSimone, Giovanni Noselli

**Affiliations:** SISSA–International School for Advanced Studies, 34136 Trieste, Italy; The BioRobotics Institute, Scuola Superiore Sant’Anna, 56127 Pisa, Italy

**Keywords:** plant morphogenesis, 3D morphoelastic rods, differential growth, circumnutation, Hopf bifurcation, flutter instability, two-oscillator hypothesis

## Abstract

We present a three-dimensional morphoelastic rod model capable to describe the morphogenesis of growing plant shoots, as driven by differential growth at the tip. We discuss the evolution laws for endogenous oscillators, straightening mechanisms and reorientations to directional cues, such as phototropic responses to a far light source and gravitropic reactions governed by the statoliths avalanche dynamics. We use this model to investigate the role of elastic deflections due to gravity loading in circumnutating plant shoots. We show that, in the absence of endogenous cues, pendular and circular oscillations arise as a critical length is attained, thus suggesting the occurrence of a Hopf bifurcation reminiscent of flutter instabilities exhibited by structural systems under nonconservative loads. When also oscillations due to endogenous cues are present, their weight relative to those associated with the Hopf instability varies in time as the shoot length and other biomechanical properties change. Thanks to the simultaneous occurrence of these two oscillatory mechanisms, we are able to reproduce a variety of complex behaviors, including trochoid-like patterns, which evolve into circular orbits as the shoot length increases, and the amplitude of the flutter induced oscillations becomes dominant. Our findings suggest that the relative importance of the two mechanisms is an emergent property of the system that is affected by the amplitude of elastic deformations, and highlight the crucial role of elasticity in the analysis of circumnutations.

## 1 Introduction

The extraordinary variety of movements in plants has fascinated scientists since the pioneering work by Darwin [11], and is raising considerable and growing interest. Many essential functions, such as reproduction, nutrition and defense, involve passive conformational changes and active adaptation triggered by diverse conditions. Indeed tropic responses and nutational movements, explosive seed and pollen dispersal, and phenomena such as the snapping of *Venus flytrap* or the closing of *Mimosa Pudica*, provide spectacular illustrations of how active biochemical processes and mechanical instabilities cooperate in plant architectures in order to produce a function [13, 18]. The principles and methods of mechanics have been successfully extended and applied to obtain biological insight into many of these plant behaviours, to investigate hypotheses and validate theories. In the context of development and morphogenesis of slender plant organs, significant advances in the modeling of plant response to a variety of cues (*e.g.*, gravity, bending and contact) have been obtained in the last decades. Nevertheless, results on the way complex three-dimensional dynamics of growing organs is affected by elastic deformations are still very limited.

Here we propose a three-dimensional model for growing plant shoots building upon the theory of morphoelastic rods. This provides a general framework to model elongating slender structures in space by efficiently decoupling growth and remodeling processes from mechanical and elastic deformations [14]. This is done by introducing an unstressed virtual configuration, where the rod is free to grow and evolve in the absence of loads and boundary conditions. The distinction between current and virtual configuration reflects the separation between sensing and actuating mechanisms. Plants sense the stimulus in the current configuration by means of a specific sensing apparatus, and reorient accordingly by differential growth, which provides the source term for the evolution of the virtual configuration. In this framework, we discuss the evolution laws that model the effect of internal oscillators, of reorientations under directional cues, such as gravitropic responses governed by the statoliths avalanche dynamics [9], and of straightening mechanisms as proprioceptive reactions to geometric curvatures [4]. The overall plant response results from the superposition of the response to different signals, each properly integrated in time to take delay and memory into account, as done in recent studies [9, 25].

Then we apply this model to the study of circumnutations, namely, pendular, elliptical or circular oscillatory movements in growing plant shoots such as the primary inflorescences of *Arabidopsis thaliana* illustrated in Fig. 1, see also supplemental video 1. The nature of these phenomena has been intensively investigated over the last century, and this produced three main hypotheses. First, as already suggested by Darwin [11], oscillatory movements might be driven by endogenous oscillators, internally regulating differential growth. Second, circumnutations might be the byproduct of posture control mechanisms that overshoot the target equilibrium, due to delayed responses [15]. Third, the previous two mechanisms might be combined in a “two-oscillator” hypothesis in which endogenous prescriptions and delayed responses coexist [19]. The overshooting hypothesis is typically based on externally driven feedback systems (of gravitropic, autotropic, phototropic or other nature) and mechanical (elastic) deformations of the plant organ are neglected. In this way, mechanical parameters play no role in controlling the occurrence of exogenous oscillations. However, as shown by the authors in a recent study [1], accounting for elastic deflections due to gravity loading enriches the scenario with spontaneous oscillations that might arise as system instabilities (bifurcations) when a loading parameter exceeds a critical value. We refer to this revised scenario as the “mechanical flutter” hypothesis, as circumnutations are reminiscent of dynamic instabilities exhibited by mechanical system under nonconservative loads [7, 6]. We extend the study presented by Agostinelli et al. [1] to the three-dimensional case by including the effect of inherent straightening mechanisms (proprioception).

**Figure 1:**
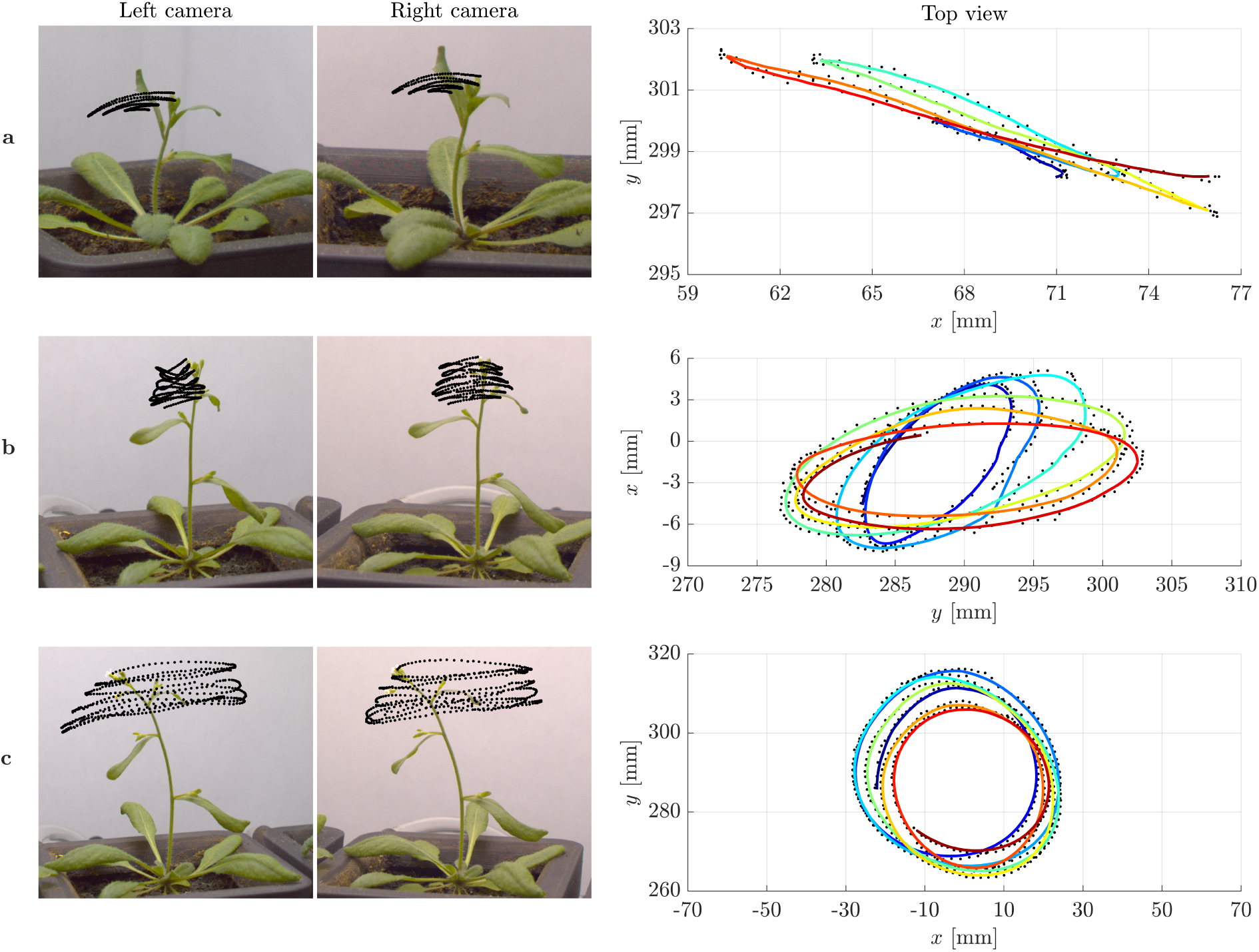
Examples of tip trajectories from specimens of *Arabidopsis thaliana* (ecotype Col-0) grown under normal gravity conditions (1 g) and continuous light at the SAMBA laboratory of SISSA: (a) Pendular oscillations in specimen 1 (about 27 days old), (b) elliptic and (c) circular patterns in specimen 2 (about 29 days old). Left: Stereo pair of images corresponding to the last instant of the tip trajectories. The superimposed black dots are the tracked positions of the tip at time intervals of 1 minute. Right: Top view of the tip trajectories as reconstructed by matching corresponding points in the stereo pair of images. The colored lines, from blue to red for increasing time, are obtained by moving averaging over ten detected positions, shown in black. Notice that the characteristic time of circumnutational oscillations *τ*_*c*_ is of the order of 70-90 min.

As discussed in the paper, we calibrate the model in agreement with results from the relevant literature and we identify the regime of model parameters for which the bifurcation is likely to occur. We find that, in the presence of gravity, the bifurcation is associated with the shoot length exceeding a critical value *ℓ*^⋆^ that depends on several parameters. These include the growth time *τ*_*g*_, the characteristic times of gravitropism and proprioception, namely, the memory times *τ*_*m*_ and 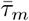, and the reaction times *τ*_*r*_ and 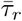, as well as morphological and biomechanical parameters, such as the organ radius *r*, the mass density *ρ*, and the stiffnesses *K*_*j*_. The presence of a proprioceptive term has an influence on the value of *ℓ*^⋆^ but it does not hinder the bifurcation phenomenon. In addition, proprioception may induce spontaneous oscillations when the growth rate exceeds a critical value 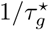, which is independent of the shoot length. As already proposed in previous studies, this feature may provide an explanation for oscillations in microgravity conditions [19]. However, the value of 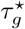 that we find with our model calibration turns out to be one order of magnitude smaller than reported experimental observations. Finally, in the presence of oscillations of endogenous origin, their relative importance with respect to the ones associated with the flutter mechanism varies in time as the biomechanical properties and the shoot length change. When all the parameters but the shoot length are fixed, elastic deformations due to gravity loading become increasingly important as the plant organ grows, and the oscillations associated with the flutter mechanism become dominant over those of endogenous origin, see supplemental videos 2 and 3. In intermediate regimes, we find trochoid-like patterns that are reminiscent of the trajectories observed by Schuster and Engelmann [33] in the hypocotyls of *Arabidopsis thaliana*.

## 2 A 3D morphoelastic rod model for growing slender organs

Building on the theory of morphoelasticity, we propose in this section a 3D rod model to describe elongating, slender plant organs. We do so under the key assumption that the time-scale for mechanical equilibrium is much shorter than any biological time-scale of the plant.

### 2.1 Kinematics: Initial, virtual and current configuration

Consider an Euclidean space 𝔼^3^ with a fixed right-handed orthonormal basis {**e**_1_, **e**_2_, **e**_3_} and define three different configurations of the rod:

- An initial reference configuration *ℬ*_0_ given by a rod having axis **p**_0_(*S*) and material cross sections characterized by the orthonormal directors 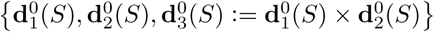 where *S* ∈ [0, *ℓ*_0_] is the arc length material parameter describing the distance from the base;
- A virtual reference configuration *ℬ*_*v*_(*t*) defined as the unstressed realization of the rod at time *t*, having axis **p**_*v*_(*s*_*v*_, *t*) and orthonormal directors 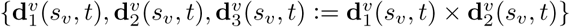 where *s*_*v*_ ∈ [0, *ℓ*_*v*_(*t*)] is the arc length coordinate;
- A current configuration *ℬ* (*t*) which is the actual shape of the rod at time *t*, taking into account deflection from mechanical loads and boundary conditions. Such a rod is defined by the space curve **p**(*s, t*) equipped with the triple of right-handed orthonormal directors {**d**_1_(*s, t*), **d**_2_(*s, t*), **d**_3_(*s, t*) := **d**_1_(*s, t*) *×* **d**_2_(*s, t*)} where *s* ∈ [0, *ℓ* (*t*)] is the arc length parameter.

In particular, we choose the initial reference configuration as the virtual configuration at time *t* = 0, namely, *ℓ*_0_ := *ℓ*_*v*_(0) and 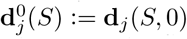 for all *S* ∈ [0, *ℓ*_0_].

Since the parameter *S* is a material coordinate for both the virtual and the current configuration, we define the respective motions, namely,

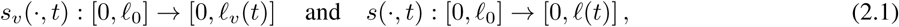

and we denote their inverse functions by the same symbol *S*(·, *t*). Moreover, in order to simplify the notation, we use the same symbol to denote material and spatial descriptions of any given field. Therefore any field defined on one of the three configurations can be evaluated at each of the other ones, by means of an implicit composition of functions. For example, given a Lagrangian (or material) field *f* (*S, t*) : [0, *ℓ*_0_] → ℝ, the associated Eulerian (or spatial) field is simply denoted by *f* (*s, t*) := *f* (*S*(*s, t*), *t*) : [0, *ℓ* (*t*)] → ℝ.

In the following we use a superimposed dot to denote the material time derivatives of any spatial vector or scalar field. In this framework we introduce the *full axial stretch* as

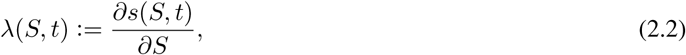

which can be decomposed in the product *λ*(*S, t*) = *σ*(*s*_*v*_(*S, t*), *t*)*γ*(*S, t*) where

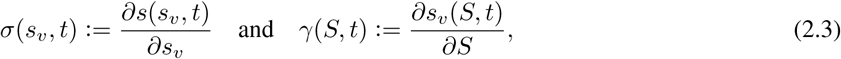

are the *elastic stretch* and the *growth stretch*, respectively. Then, we define the *true strains*

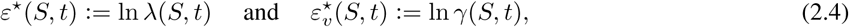

which will turn out to be crucial growth quantifier.

From classical rod theory [2], we know that there exist vector-valued functions **u**(*s, t*), called *twist*, and **w**(*s, t*), called *spin*, such that

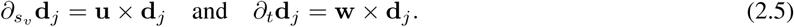

As for the components *u*_*j*_ := **u** · **d**_*j*_, these are referred to as *flexural strains* for *j* = 1, 2 and *torsional strain* for *j* = 3. In a similar manner, the directors of the virtual configuration define the *spontaneous twist*, **u**^⋆^, and the *spontaneous spin*, **w**^⋆^, *i.e.*,

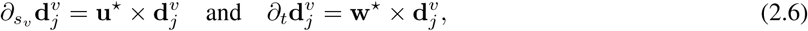

and the components 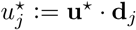 are called *spontaneous strains*.

### 2.2 Mechanics and constitutive assumptions

Under the quasi-static assumption we impose the static equilibrium in the virtual reference configuration at all times, such that

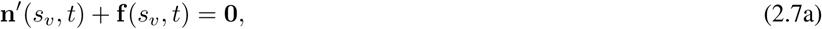

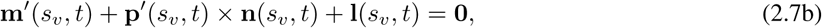

where a prime denotes differentiation with respect to *s*_*v*_, **n** and **m** are the resultant contact force and contact couple, whereas **f** and **l** are the body force and couple per unit virtual reference length, respectively. Determination of the current configuration *ℬ* (*t*) can be achieved by solving equations (2.7) combined with a suitable constitutive model and appropriate boundary conditions.

For plant organs, a reasonable assumption is to treat them as unshearable (**d**_3_ = ∂_*s*_**p**) and elastically inextensible rods, such that *σ* = 1 and *s* = *s*_*v*_, and characterized by a quadratic strain-energy function defined by a diagonal stiffness matrix **K**, such that the constitutive law reads

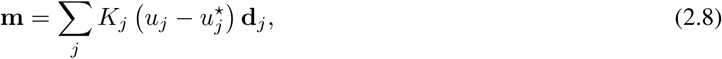

where *K*_*j*_ denotes the *j*-th diagonal component of **K**. More specifically, in this study we assume rods of circular cross section of radius *r*, such that *K*_1_ = *K*_2_ = *EI*, where *E* is the Young’s modulus and *I* = *πr*^4^*/*4 is the second moment of inertia, and *K*_3_ = *µJ* where *J* = 2*I* and *µ* = 2*E*(1 + *ν*) is the shear modulus determined by the Poisson’s ratio *ν*. In passing, we notice that such a modeling assumption might be refined by considering elliptic cross sections, which provide more accurate descriptions of some plant organs [30].

In the absence of external loads and couples, *i.e.*, for **f** = **0** and **l** = **0**, equations (2.7) lead to **n** = **0** and **m** = **0**, in which case the visible strains coincide with the spontaneous strains of the virtual configuration, *ℬ*_*v*_(*t*). Therefore, in this special case we recover the kinematic model recently proposed by Porat et al. [32].

### 2.3 Tip growth

In both roots and shoots, tip (or primary) growth can be modeled as a process localized at the end of the organ, in a region of constant size *ℓ*_*g*_. As for the growth stretch *γ* and the true strain 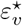, these are two connected key quantifiers in the modeling of growth by elongation. Indeed, the *relative elemental growth rate* (REGR) introduced by Erickson and Sax [12] is precisely given by the strain rate 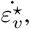, which can be written in terms of the growth stretch as

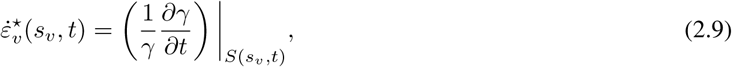

where a dot denotes the material time derivative, cf. Supplemental Section S1. Since such a quantity can be experimentally measured by tracking material markers along the organ [23, 5, 26, 36, 17, 31], we prescribe a function *G*(*s*_*v*_, *t*), vanishing outside the elongating zone [*ℓ*_*v*_(*t*) − *ℓ*_*g*_, *ℓ*_*v*_(*t*)] and such that 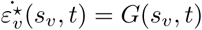. Consequently, tip growth is governed by two coupled PDEs, namely,

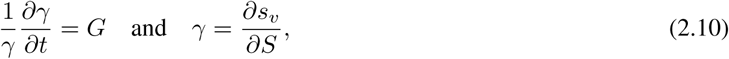

to be solved for *S* ∈ [0, *ℓ*_0_] and *t* ≥ 0, with initial condition *γ*(·, 0) ≡ 1 and fixed boundary datum *s*_*v*_(0, ·) ≡ 0. We report in Fig. 2 some solutions of problem (2.10) for REGR profiles of the form

**Figure 2:**
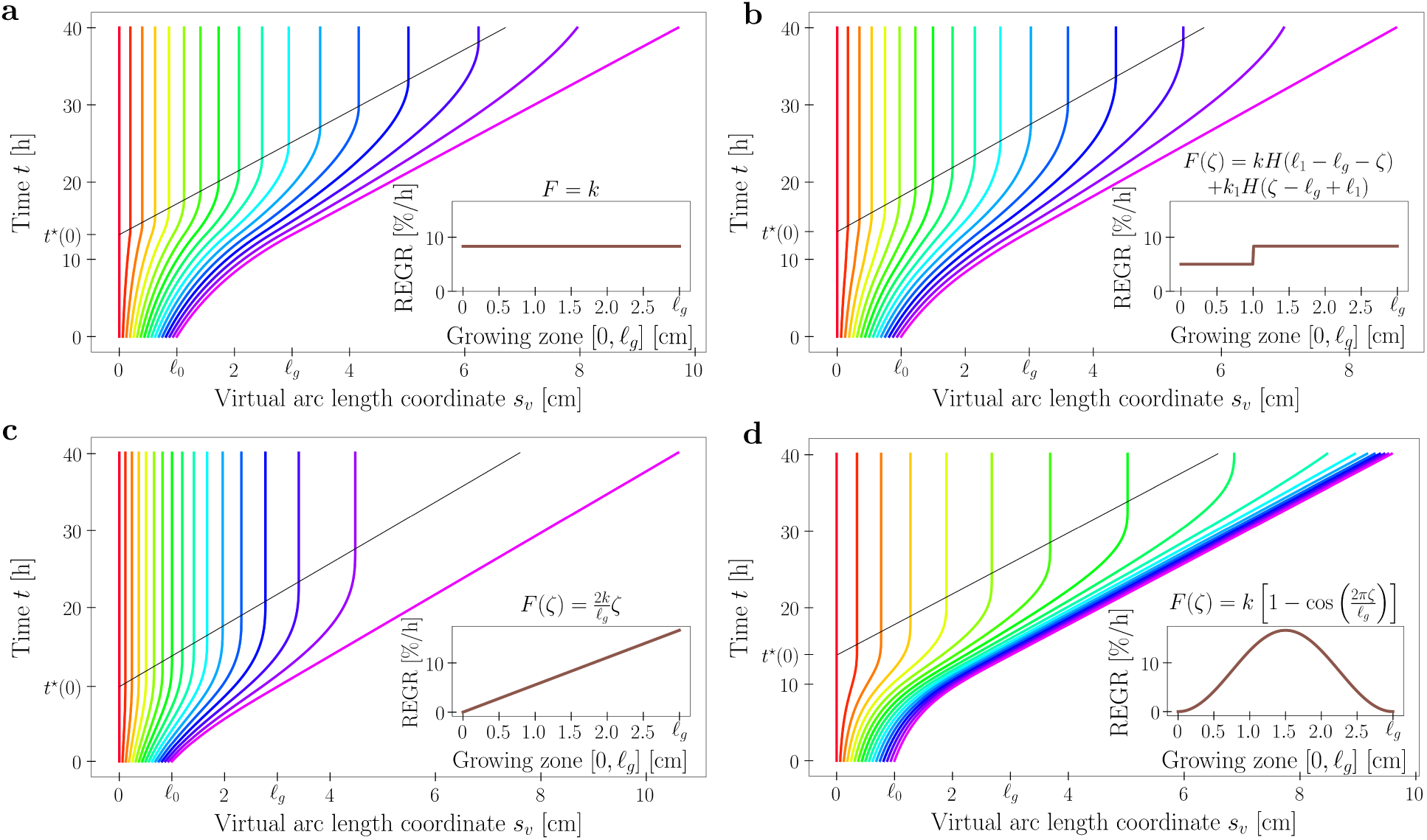
Time evolution of the virtual arc length *s*_*v*_ (*S, t*) for a set of 17 material points along the plant organ for different functions *G* of the form (2.11): (a) *F* (*ζ*) = *k*, (b) *F* (*ζ*) = *kH*(*ℓ*_1_ − *ℓ*_*g*_ − *ζ*) + *k*_1_*H*(*ζ* − *ℓ*_*g*_ + − *ℓ*_1_), (c) *F* (*ζ*) = 2*kζ/ ℓ*_*g*_, (d) *F* (*ζ*) = *k* (1 − cos (2*πζ/ ℓ*_*g*_)). Model parameters are *ℓ*_0_ = 1 cm, *ℓ*_*g*_ = 3 cm, *ℓ*_1_ = 2 cm, *k* = 0.05 h^−1^ and *k*_1_ = 0.083 h^−1^. Notice the black line that denotes the time *t*^∗^(*S*) at which the material point *S* exits the growth zone, as its distance from the tip exceeds *ℓ*_*g*_. We refer to Supplemental Section S1 for additional details.

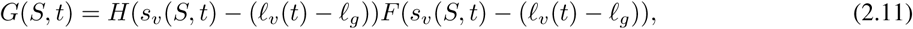

where *H*(·) is the Heaviside function and *F* : [0, *ℓ*_*g*_] → ℝ_+_ is a nonzero continuous function. In this case, for *ℓ*_0_ ≥ *ℓ*_*g*_ we get

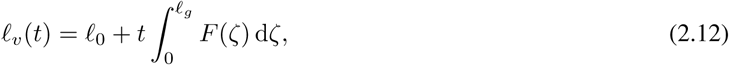

which is a linear function of time regardless of the particular choice of *F*. We refer to Supplemental Section S1 for a more detailed derivation of equation (2.9) and for analytical and computational solutions of problem (2.10).

### 2.4 Differential growth and evolution laws

The shape of growing plant roots and shoots evolves and adapts by responding to a variety of stimuli. The main morphing mechanism consists in a spatially nonhomogeneous growth rate of the cross section, called *differential growth*. For any cross section *s*_*v*_ and time *t*, equation (2.9) can be used to extend the notion of relative elemental growth rate to any point (*x, y*) of the cross section: We denote it by 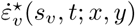, cf. Supplemental Section S2. Then, by a Taylor expansion about the center of the cross section we arrive at

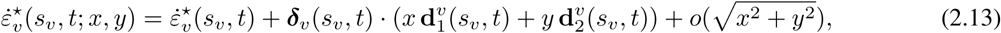

where *x* and *y* are the coordinates of the point in the local basis 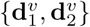, and

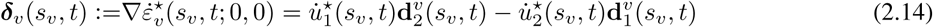

is the *growth gradient* on the virtual cross section *s*_*v*_ at time *t*. Hence the corresponding growth gradient in the current configuration is given by

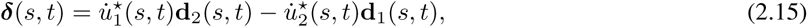

which is orthogonal to the axis of bending, see Fig. 3a.

**Figure 3:**
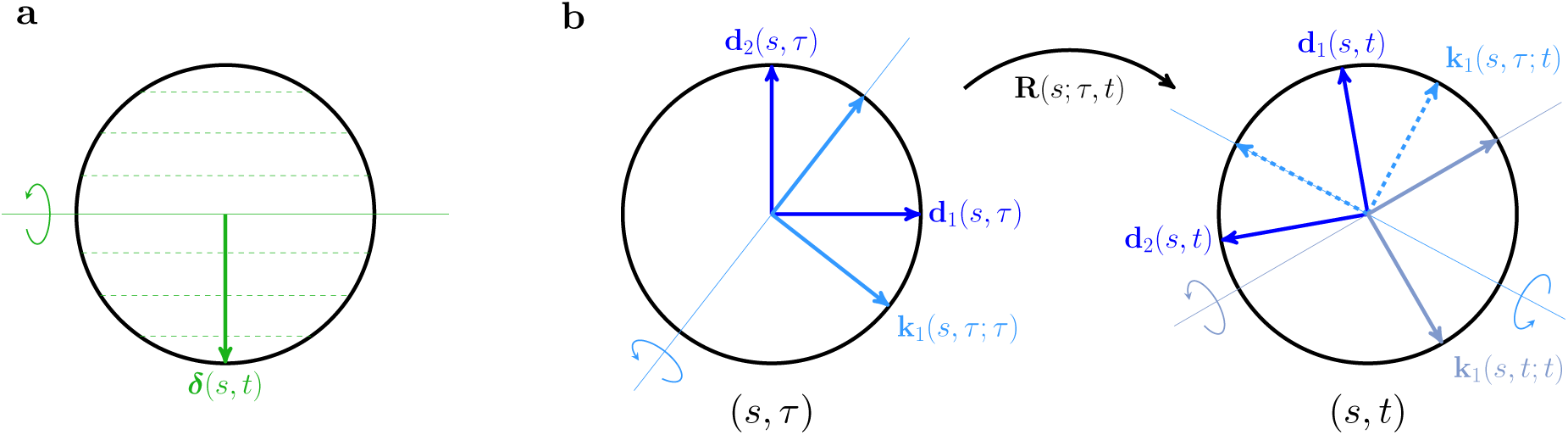
(a) Level curves of an affine strain rate having gradient ***δ***, on the cross section (*s, t*); the axis perpendicular to ***δ*** is the one about which bending occurs as due to differential growth. (b) Time evolution of the material cross section. **R**(*s*; *τ, t*) is the rotation mapping **d**_*j*_ (*s, τ*) into **d**_*j*_ (*s, t*), whereas **k**_1_(*s, τ* ; *t*) denotes the contribution to the growth gradient at time *t* due to a stimulus sensed at time *τ*.

Equation (2.15) reveals the connection between differential growth and spontaneous strain rates. Indeed when the growth rate of the cross section is affine, or the organ radius is small enough to justify a linearization, the prescription of the growth gradient ***δ*** results in the evolution laws for the spontaneous flexural strains 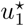 and 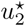. We observe that the contribution of the torsional strain 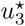 to the growth gradient is negligible. Nevertheless, it could play a crucial role in other growth mechanisms, such as that observed in twining plants.

In the presence of *n* different stimuli, we assume a weighted average of their respective growth gradients defined on the current cross section. In other terms, the overall growth gradient for a circular cross section of radius *r* is determined by

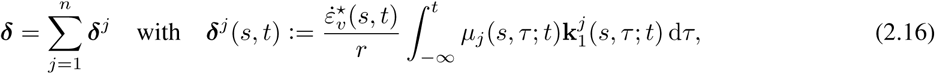

where 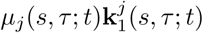 is the vector on the *current* cross section that defines the contribution to the growth gradient from the *j*-th stimulus sensed at time *τ*. Equation (2.16) can be projected on the local basis {**d**_1_, **d**_2_} to get

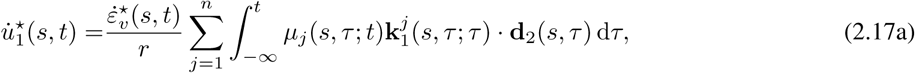

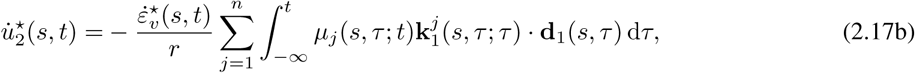

where we have used the fact that the contribution to growth sensed at a certain time *τ* is fixed in the frame of the directors, namely, **k**_1_(*s, τ* ; *t*) = **R**(*s*; *τ, t*)**k**_1_(*s, τ* ; *τ*) where **R**(*s*; *τ, t*) is the rotation that maps **d**_*j*_(*s, τ*) into **d**_*j*_(*s, t*), as illustrated in Fig. 3b.

We use equations (2.17) to describe a response of a material cross section to a stimulus sensed at the very same location. However, they might be adapted to the case of nonlocal responses, as it occurs for gravitropic reactions of roots [27]. Moreover, these expressions allow to include memory and delay effects, as done in recent studies [9, 25], and the instantaneous models are recovered as special cases by choosing the Dirac delta as response function.

In the following we discuss the plant response to different stimuli: Endogenous prescription (*e.g*, internal oscillators), reorientation to align the organ axis with a given vector (*e.g.*, gravitropism or phototropism), and straightening mechanism (*i.e.*, proprioception).

#### Endogenous cues

Inspired by the Darwinian concept of internal oscillator, we implement an endogenous driver for the differential growth mechanism. Several studies on plant growth and nutations have revealed a strong correlation between oscillatory movements and biological rhythms [5, 35, 34, 33, 8, 28] and three-dimensional models including endogenous mechanisms have already been proposed [4, 32]. In our framework, we extend such approaches by assigning a growth direction in the stem cross section *s* at time *t*, namely,

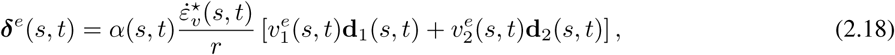

where *α* is a scalar dimensionless sensitivity parameter and 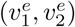 are the prescribed growth components in the local basis. As an example, we consider a spatially uniform time-harmonic oscillator, such that

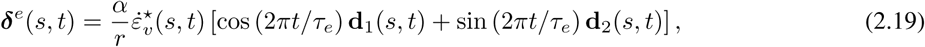

where *τ*_*e*_ is the period of endogenous oscillations and *α* is constant.

#### Reorientation under directional cues

Any vector stimulus **s** sensed in the current configuration, towards (or away from) which the plant organ aligns via differential growth (*e.g.*, gravitropism and phototropism for a far light source), contributes to growth gradient via its projection on the plane (**d**_1_, **d**_2_), such that

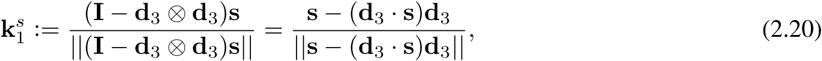

where **I** denotes the identity operator. Notice that the direction of null differential growth, about which the organ bends, is given by 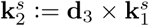. Then the growth gradient associated with the stimulus **s** can be written as

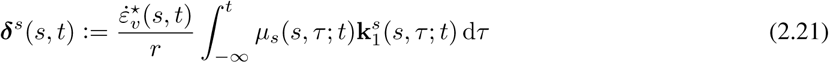

for an appropriate choice of the response function *µ*_*s*_.

When applying equation (2.20) to gravitropism, we have two choices for the stimulus **s**, which are illustrated in Fig. 4c. One possibility is to approximate the stimulus with the vector of gravitational acceleration **g**, thus neglecting the microscopic description of how gravity is sensed by plant organs. On the contrary, a more accurate choice is to consider the stimulus as perceived by the gravity sensing apparatus. In particular, it is widely accepted that plant organs sense gravity through the sedimentation of starch-filled plastids, called statoliths, into specialized cells, called statocytes, which are found along the shoot growing zone and in the root caps [9, 27]. In this case, we can assume the stimulus **s** to be given by the average outer normal to the free surface of the pile of statoliths, as shown in Fig. 4c. More specifically, by extending the approach taken by Chauvet et al. [9] to the three-dimensional case, we can model the statoliths free surface as a plane with normal **h**, whose dynamics is a viscous relaxation to − **g**. As shown in Supplemental Section S3, the time evolution of **h** is governed by

**Figure 4:**
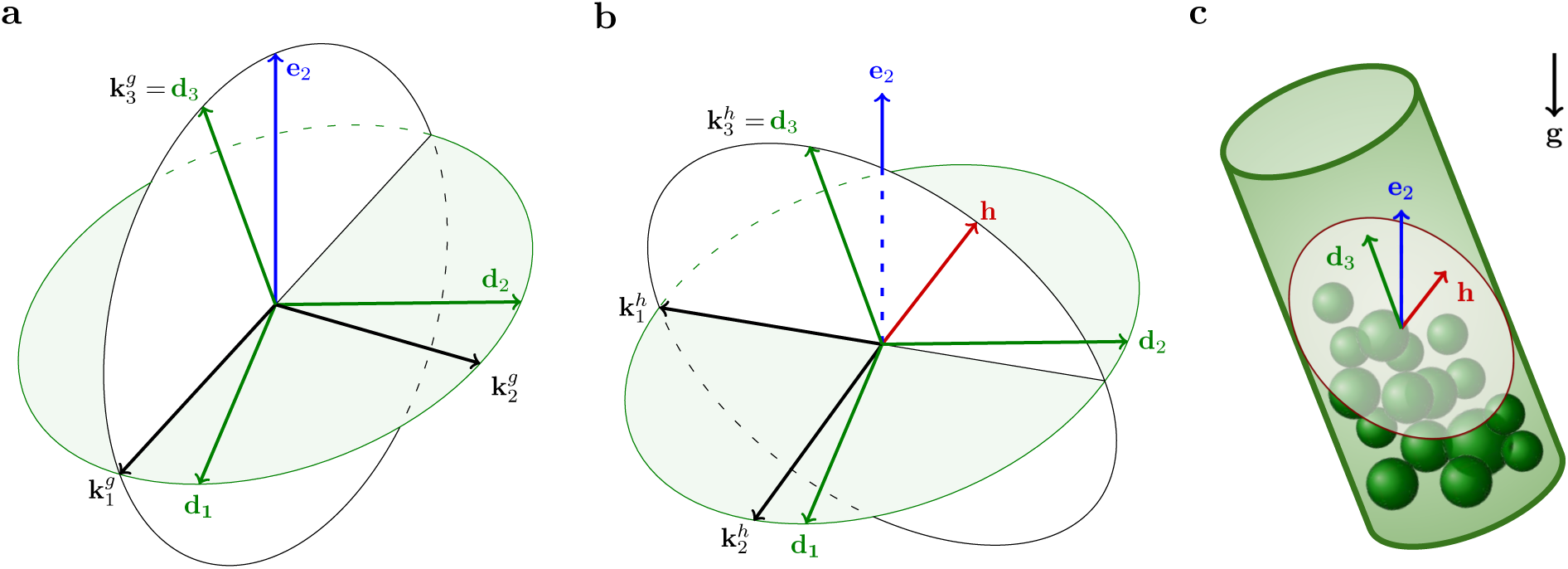
(a-b) Illustration of the orthonormal bases exploited to define the gravitropic responses. (a) The basis 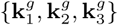 is constructed by defining 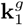 as the unit vector laying on the stem cross section and having the most negative **e**_**2**_-component, and setting 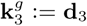 and 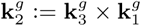. (b) The basis 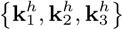 is constructed in a similar manner by defining the unit vector laying on the stem cross section having the most negative **h**-component, 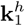, and considering 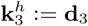 and 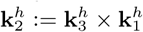. (c) Sketch of a single statocyte cell where **h** is the average outer normal to the free surface of the pile of statoliths.

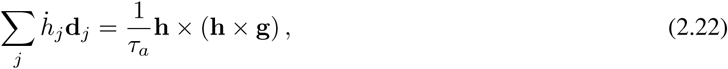

where *τ*_*a*_ is the characteristic time for the statoliths avalanche dynamics and *h*_*j*_ := **h** · **d**_*j*_. As elsewhere, a superimposed dot denotes the material time derivative.

#### Proprioception

A number of experimental studies have pointed out the existence of an independent straightening mechanism, often referred to as proprioception, autotropism or autostraightening, which is triggered by bending of the organ [29]. Following Bastien et al. [3] and Bastien and Meroz [4], we assume that such a straightening response is driven by the geometric curvature of the organ, *i.e.*, 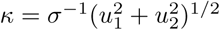, thus producing a growth gradient parallel to the visible normal vector ***ν*** := *κ*^−1^∂_*s*_**t**, where **t** = ∂_*s*_**p** is the tangent to the rod axis. In other terms, we prescribe

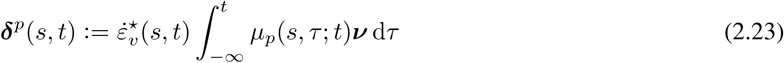

for an appropriate response function *µ*_*p*_ that is proportional to *κ*.

### 2.5 Remodeling of other parameters

Plant growth is typically accompanied by lignification processes that determine the evolution of the mechanical properties of the organ. In the current framework, this can be accounted for by rod stiffening, adapting the approach taken by Chelakkot and Mahadevan [10]; in particular, we assume the Young’s modulus to evolve in time according to

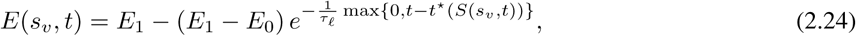

where *t*^⋆^(*S*) is the time at which the material point *S* exits the growth zone, *τ*_*ℓ*_ is the lignification characteristic time, whereas *E*_0_ and *E*_1_ are the minimum and maximum values of the Young’s modulus, respectively.

### 2.6 A model for growing plant shoots

In concluding this section, we introduce a working model for elongating plant shoots suitable for the study of circumnutations by specializing some of the mechanisms described above. We make the following assumptions.

i. As previously stated, the plant shoot is modeled as an unshearable and elastically inextensible rod of circular cross section such that the constitutive law for the moments is given by (2.8).
ii. Following previous results [3, 4, 10, 9], we specialize the tip growth model of (2.10) by choosing a piecewise constant growth function, namely,

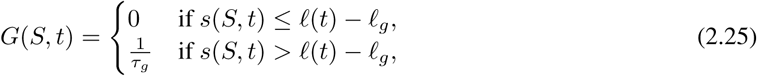

where *τ*_*g*_ > 0 is the characteristic growth time. For such a case, the map *s*(*S, t*) can be analytically determined, as shown in Supplemental Section S1.
iii. We assume that there is no external couple and that the shoot carries a uniform distributed gravity load *q* = *ρgA* ≥ 0, where *ρ* is the mass density, *A* = *πr*^2^ is the cross-sectional area, and *g* the gravitational acceleration so that **f** = *q* **g** = −*q* **e**_2_. Moreover, the apical end is free and the boundary conditions associated with equations (2.7) read **n**(*ℓ* (*t*), *t*) = **m**(*ℓ* (*t*), *t*) = **0**. Then, equation (2.7a) can be integrated to get **n** = *q* (*ℓ* (*t*) − *s*) **e**_2_ and we are left with **m**′(*s, t*) = −*q* (*ℓ* (*t*) − *s*) **e**_2_ *×* **d**_3_(*s, t*).
iv. Differential growth is due to the combination of the harmonic intrinsic oscillator (2.19) and gravitropic-proprioceptive reactions weighted by an exponential response function so that the evolution of spontaneous strains follows

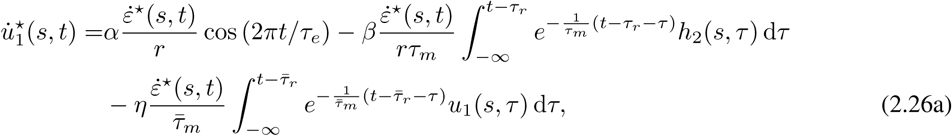

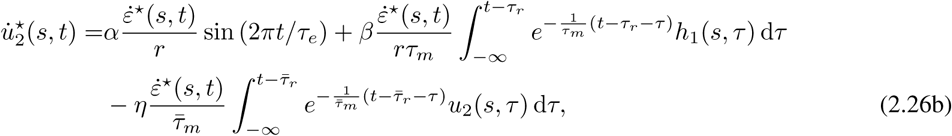

supplemented by the statoliths avalanche dynamics of (2.22). Here primes denote differentiation with respect to the parameter *s* and dots denote material time derivatives. As for the parameters *α, β, η*, these are nonnegative dimensionless sensitivities associated with the endogenous cues, gravi-responses and proprioception, respectively, whereas *τ*_*m*_ and *τ*_*r*_ (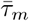 and 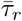) are the memory and reaction times of the gravitational (straightening) stimulus.
v. We assume that there is no evolution of the spontaneous torsional strain, namely, we set 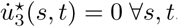.
vi. The shoot lignification process is taken into account by the remodeling of the rod stiffness given by (2.24).
vii. As for the analysis reported in the following section, the list of relevant parameters is summarized in Table 1.

## 3 Results on circumnutations

In the following we perform a study of plant circumnutations based on the proposed rod model, by gradually including more and more terms. We first explore the role of gravitropic and proprioceptive responses independently one from the other. Then we analyze the interaction of these two adaptive growth processes to understand their combined effect upon the system. Finally, we include in our analysis oscillations of endogenous nature to investigate their relative importance with respect to the oscillations caused by flutter instability, as the organ evolves in time.

**Table 1:** Summary of model parameters and respective order of magnitude of their values.

| Parameter | Description | Value | Source |
| --- | --- | --- | --- |
| Sensitivities for differential growth |  |  |  |
| $\alpha$ | sensitivity to endogenous cues | 0 – 1 | assumed |
| $\beta$ | gravitropic sensitivity | 0.8 | [9] |
| $\eta$ | proprioceptive sensitivity | 20 | assumed |
| Characteristic times |  |  |  |
| $\tau_a$ | time for statoliths avalanche | 2 min | [9] |
| $\tau_e$ | period of endogenous oscillations | 20 min | assumed |
| $\tau_m$ | gravitropic memory time | 12 min | [9] |
| $\bar{\tau}_m$ | proprioceptive memory time | 12 min | assumed |
| $\tau_r$ | gravitropic reaction time | 12 min | [9] |
| $\bar{\tau}_r$ | proprioceptive reaction time | 12 min | assumed |
| $\tau_g$ | growth time | 20 – 40 h | [9, 31, 17] |
| $\tau_\ell$ | lignification time | 6 d | assumed |
| Morphological and biomechanical parameters |  |  |  |
| $r$ | radius of the cross section | 0.5 mm | [30] |
| $\ell_g$ | growth zone | 4 – 7 cm | [17] |
| $\nu$ | Poisson's ratio | 0.5 | assumed |
| $\rho$ | mass density | $10^3 \text{ Kg m}^{-3}$ | [10] |
| $E_0$ | initial Young's modulus | 10 MPa | [10] |
| $E_1/E_0$ | stiffening ratio due to lignification | 200 | assumed |

### 3.1 The regime of short times

Under suitable assumptions on the relevant time scales we deduce a model that is amenable for theoretical analysis. More specifically, we exploit the fact that the statolith avalanche dynamics is much faster than that of circumnutations with characteristic time *τ*_*c*_, which, in turn, is much faster than the processes of organ elongation and lignification. Based on the assumption of *τ*_*g*_ ≫ *τ*_*c*_, we neglect changes in length, cf. equation (2.25), and we assume that the whole organ is “active”, *i.e., ℓ* ≈ *ℓ*_0_ ≤ *ℓ*_*g*_. In this case, current and reference arc lengths coincide such that material time derivatives reduce to standard time derivatives. As for the condition on the lignification time, *i.e., τ*_*ℓ*_ ≫ *τ*_*c*_, this implies that at short times compared to *τ*_*ℓ*_, the Young’s modulus (2.24) is approximately constant, namely *E* ≈ *E*_0_. Finally, the assumption of *τ*_*a*_ ≪*τ*_*c*_ implies **h** ≈ − **g** = **e**_2_, which is the stable steady solution to equation (2.22), so that *h*_*j*_ ≈ **d**_*j*_ · **e**_2_ for *j* = 1, 2, and the statoliths avalanche dynamics can be disregarded.

We exploit this simplified model to explore the emergence of periodic, oscillatory movements in different scenarios, by means of both theoretical and computational studies. We now summarize our findings by referring to the Supplemental Section S4 for the details of the linearized stability analyses of the trivial equilibrium, and to the Supplemental Section S5 for the description of the numerical model. The description of the Supplemental Videos is reported in the Supplemental Section S6.

#### Graviceptive case: *α* = *η* = 0 and *β* > 0

In the absence of both endogenous cues (*α* = 0) and straightening mechanisms (*η* = 0), the linearized stability analysis coincides with the one carried out for the planar model studied in [1]. We find that the straight equilibrium configuration suffers a flutter instability as the plant shoot attains a critical length. We report in Fig. 5a the stability boundary for the system in terms of the model parameters (*τ*_*g*_, *ℓ*). Moreover, we confirm by means of numerical simulations the appearance of limit cycles in the nonlinear regime and show that the planar oscillations previously reported in [1] are unstable periodic solutions, whereas three-dimensional circular patterns emerge as stable limit cycles.

**Figure 5:**
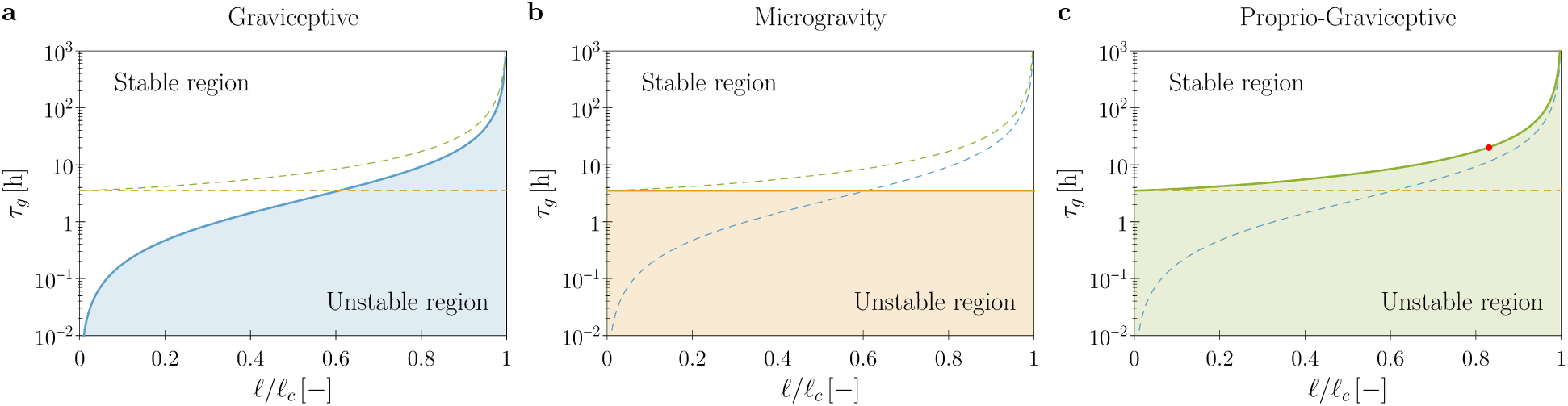
Theoretical stability boundaries in terms of the model parameters (*τ*_*g*_, *ℓ*). Blue, orange and green curves are for the graviceptive case (*α* = *η* = 0, *β* = 0.8), for microgravity (*α* = *β* = 0, *η* = 20, *q* = 0) and for the proprio-graviceptive case (*α* = 0, *β* = 0.8, *γ* = 20), respectively. In each plot (a-c) results for the relevant case are reported as solid curves, whereas the boundaries for the other two cases are shown as dashed curves for comparison purposes. Model parameters are those reported in Table 1. Shoot length is normalized by the self-buckling length 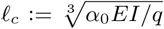 where *α*_0_ ≈ 7.837 [16]. The red dot in (c) corresponds to the computational results of Fig. 6.

#### Microgravity

*α* = *β* = 0, *η* > 0, and *q* = 0. In agreement with previous studies [19] we find that, in microgravity conditions (*β* = 0 and *q* = 0), proprioceptive responses alone might induce spontaneous oscillations. Indeed, even in the absence of an intrinsic oscillator (*α* = 0), the rest state undergoes an instability when the growth rate exceeds a critical threshold 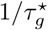. Interestingly, this is independent of the shoot length, as shown in Fig. 5b. For the model parameters of Table 1, we find a critical value of 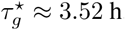 h. This seems to be out of the range of experimental observations, thus suggesting that the persistence of oscillations in microgravity might have an endogenous origin [20, 21].

#### Proprio-graviceptive case: *α* = 0 and *β, η* > 0

When proprioception and gravitropism coexist in the absence of endogenous oscillators (*α* = 0), we find the persistence of a critical growth rate 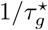, beyond which the rest state changes its stability character, as observed in microgravity. As depicted in Fig. 5c, the system may still lose stability at lower growth rates 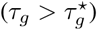 for a critical length *ℓ*^⋆^. This is different from the one found in the graviceptive case, and a numerical study of the nonlinear regime reveals the occurrence of pendular and circular limit cycles for supercritical lengths, see Fig. 6 and Supplemental Video 4. As for the effect of the auto-straightening mechanism, this lowers the critical length, provided that the delay 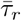 and the memory time 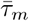 are sufficiently large.

**Figure 6:**
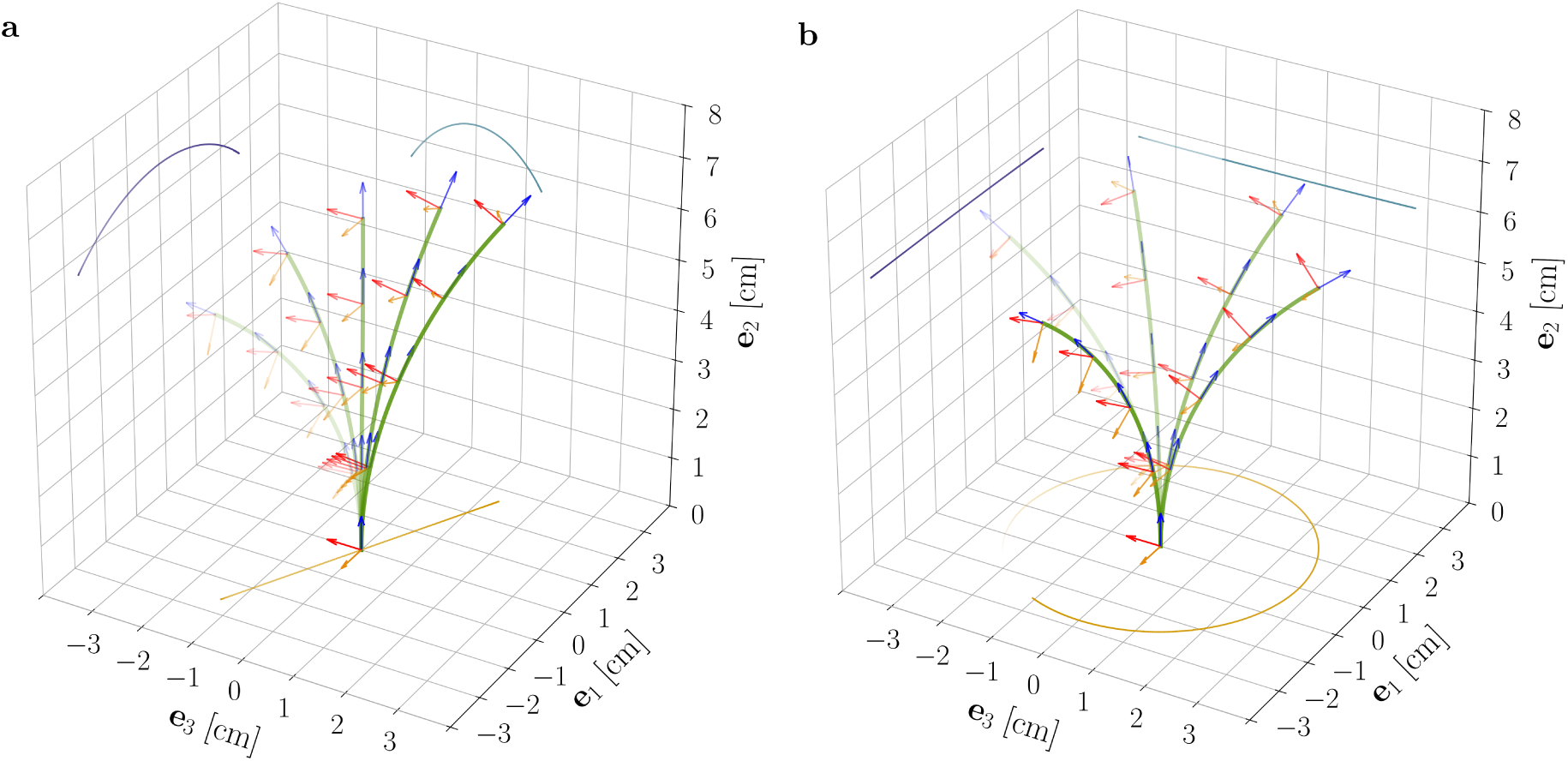
Superposition of deformed shapes and respective directors, from the reduced nonlinear rod model for *ℓ/ ℓ*_*c*_ ≈ 0.83 (= 6.59 cm), *τ*_*g*_ = 20, and the parameters reported in Table 1. This choice of model parameters corresponds to the red dot shown in Fig. 5c. For supercritical lengths (*ℓ*^⋆^ ≈ 6.56 cm, for such a choice of model parameters) two types of nontrivial periodic solutions emerge: (a) unstable pendular oscillations and (b) stable circular oscillatory patterns.

#### Endogenous oscillations

*α, β, η* > 0. By means of the computational model described in Supplemental Section S5, we consider the case of *α* > 0 to investigate the effects of an endogenous, time-harmonic oscillator with period *τ*_*e*_. For subcritical lengths, the intrinsic oscillator dominates the dynamics and the solutions ultimately converge to motions of period *τ*_*e*_. On the contrary, for supercritical lengths we find sustained dynamics for which the tip projection on the (**e**_1_, **e**_3_) plane determines trochoid-like patterns, see Fig. 7 and Supplemental Videos 5 and 6. The shape of the trochoid is determined by the ratio of two periods, namely, the one of the internal oscillator, *τ*_*e*_, and the one of the limit cycle emerging from flutter instability. As a consequence, we do not expect these patterns to be periodic unless such a ratio is a rational number. More specifically, patterns similar to epitrochoid or hypotrochoid are found when the rotational directions of the two oscillatory mechanisms are concordant or discordant, respectively. The results of Fig. 7 and the Supplemental Videos 5 and 6 exemplify the rod dynamics, together with the tip projections on the coordinate planes.

**Figure 7:**
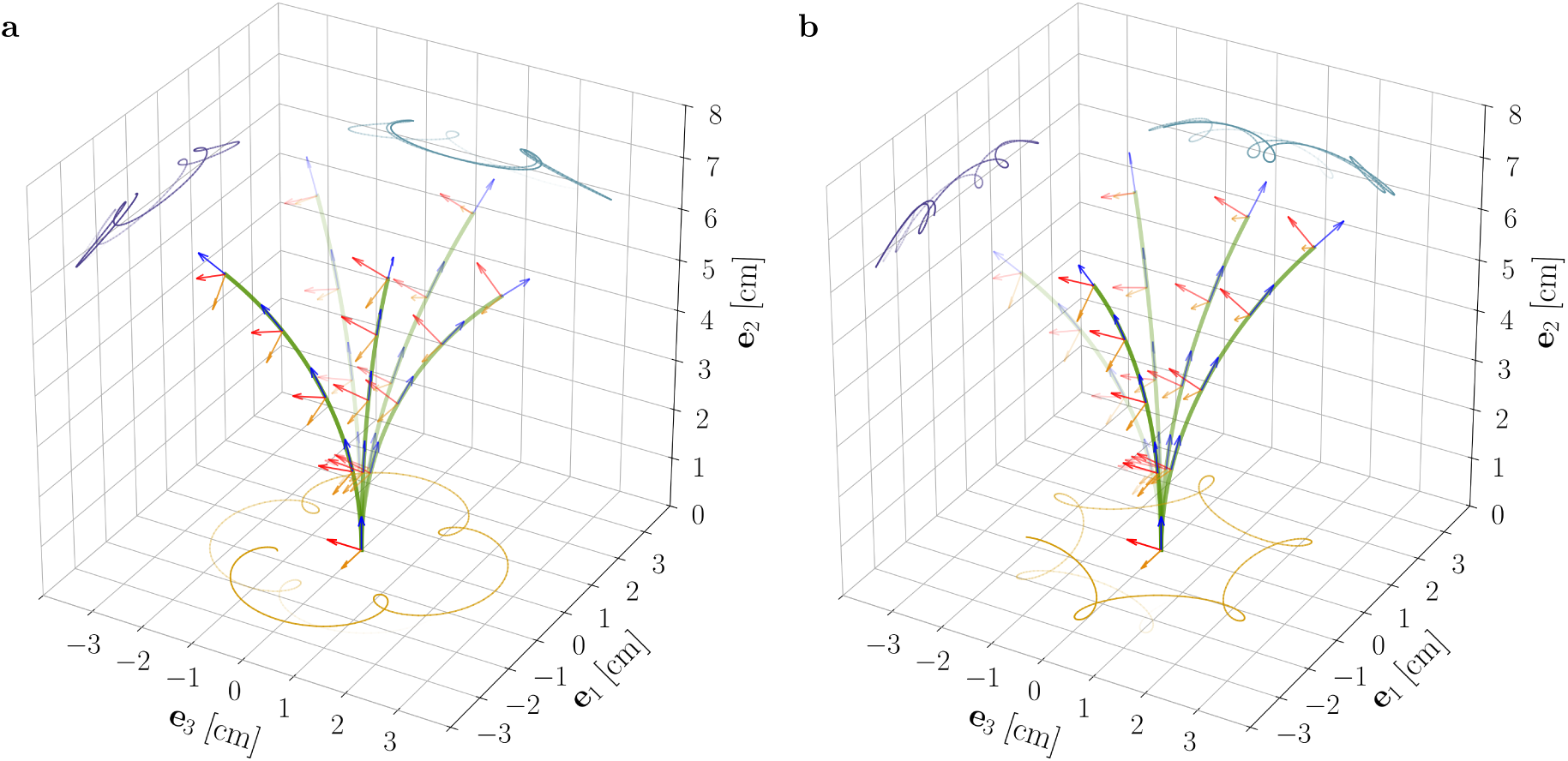
Superposition of deformed shapes and respective directors, from the reduced nonlinear rod model for *ℓ* = 6.565 cm, *α* = 0.3, and for the model parameters as reported in Table 1. Flutter was initiated in the clockwise direction by suitable initial perturbations and epitrochoid-like (a) and hypotrochoid-like (b) patterns were obtained for concordant and discordant endogenous oscillations, respectively.

### 3.2 The role of plant shoot elongation

We conclude our analysis by exploring the contribution of length changes and lignification processes in the overall dynamics of the model plant, by exploiting the computational model detailed in Supplemental Section S5. As the shoot length varies in time, the relative weight of the two oscillatory mechanisms, namely, the intrinsic oscillator and the flutter instability, changes and affects the resulting dynamics. As exemplified by Fig. 8 and the Supplemental Video 2, the system gradually transitions from a dynamics mainly characterized by endogenous oscillations in the subcritical regime (*ℓ* < *ℓ*^⋆^) to one in which flutter-induced oscillations dominate in the supercritical regime (*ℓ* > *ℓ*^⋆^). Trochoid-like patterns are visible in the intermediate regime of flutter initiation.

**Figure 8:**
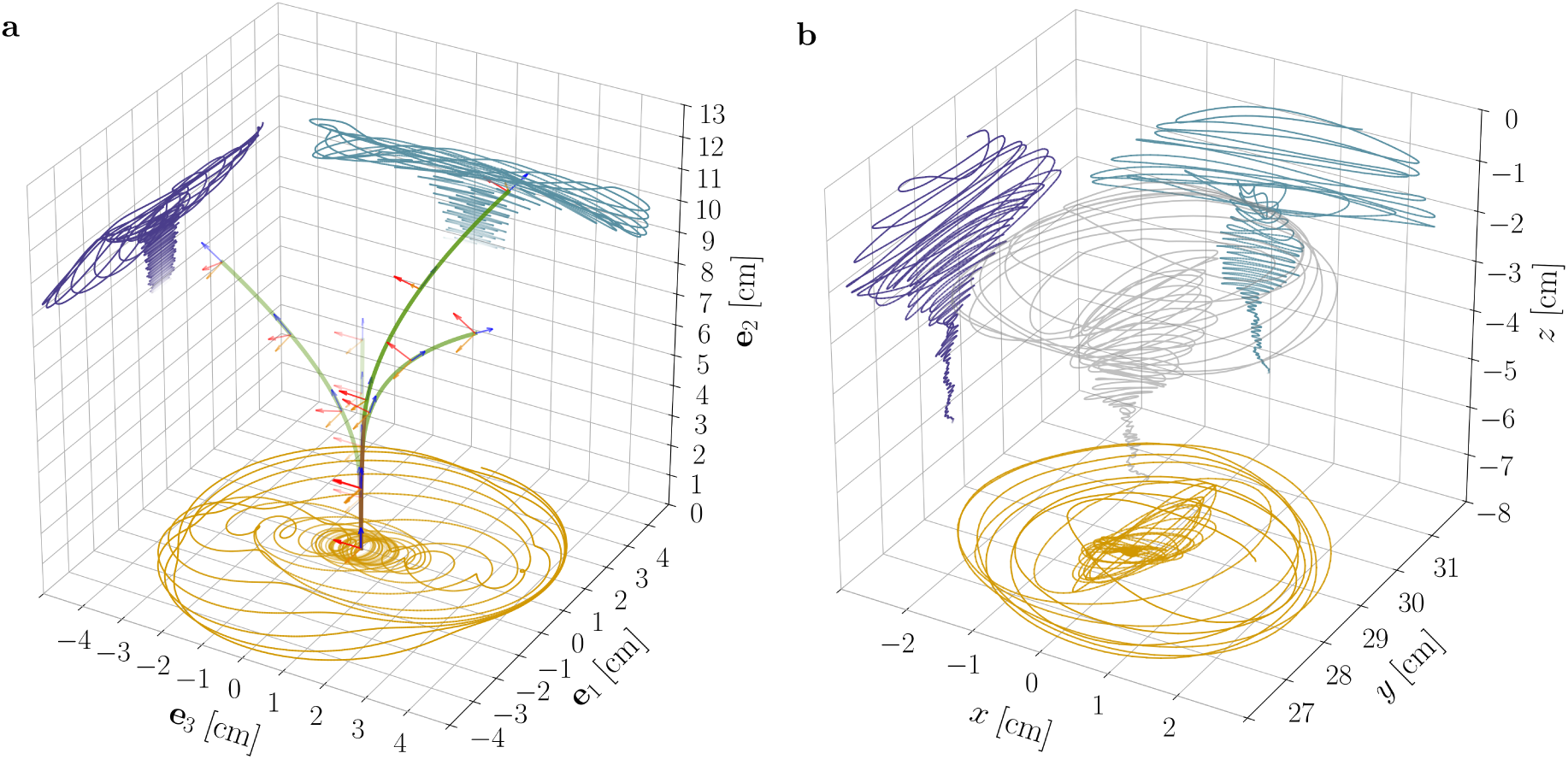
(a) Superposition of deformed shapes and respective directors from the nonlinear rod model and for the model parameters as in Supplemental Video 2. Notice the progressive transition of the system from a dynamics dominated by the endogenous oscillator to one in which flutter-induced oscillations prevail. (b) Experimental results (tip trajectory and its projections on coordinate planes) from a sample of *Arabidopsis thaliana* (Col-0) are reported for qualitative comparison, see also Supplemental Video 1.

## 4 Conclusions and outlook

Building on the general framework of morphoelasticity, we have introduced a rod model capable of describing three-dimensional motions of growing plant shoots and that accounts for directional responses driven by differential growth. These include any plant response that aligns the organ axis with a directional stimulus, *e.g.*, gravitropism and phototropism (for a far light source) as well as straightening mechanisms triggered by curvature perception. Some models previously proposed in the literature can be derived as limit cases of the present one by assigning suitable response functions and by either constraining the organ to a plane [1, 10] or disregarding elastic deformations [32] or both [3, 9, 25]. We believe that these features, namely, three-dimensionality and elasticity, play a crucial role in many phenomena and cannot be disregarded. The model proposed in this paper is intended as a test bed for different hypotheses, that may provide new biological insight. In addition, the model may be useful in the context of bioinspired soft robotics, which has recently started to draw inspiration from the plant kingdom to conceive and design innovative adaptable robots and smart-actuation strategies [24].

As an illustration of the applicability of the present model, we have studied circumnutations of growing plant shoots. Since the first experimental observations, a long-lasting debate has produced three main theories for their nature: The existence of an endogenous oscillator [11], a gravitropic feedback oscillator [15] or a combination of the two [19]. Previous analyses of these theories systematically disregarded elastic deflections due to gravity loading, which however may affect in a relevant way the mechanical stability of the biological system [1]. The present study extends these observations in various respects. First, straightening mechanisms, modeled as proprioception with delay and memory, do not alter the scenario of mechanical instabilities from a qualitative viewpoint. As proposed in the literature, in the absence of other stimuli, proprioception might even be responsible for oscillatory movements in microgravity conditions. However, for the present model calibration, the critical growth rate we determine is about ten times larger than that of available experimental observations. Second, the instability occurs at the same critical length *f*^⋆^ found in the planar case but pendular movements are unstable in the three-dimensional setting, and elliptic trajectories could represent transient oscillations towards stable circular limit cycles. Third, the flutter instability combined with an internal harmonic oscillator can reproduce trochoid-like patterns, which were observed in previous experiments on the hypocotyls of *Arabidopsis thaliana* seedlings, as the result of the superposition of short and long period nutations [33]. In the presence of elastic deformations, the relative amplitude of the two oscillations becomes time-dependent, with endogenous oscillations prevailing in the subcritical regime of short shoots (*ℓ* < *ℓ*^⋆^) and dominant fluttering in the supercritical one of long shoots (*ℓ* > *ℓ*^⋆^).

These findings suggest the possibility to reinterpret the vast existing experimental literature from a renewed perspective. Our observations conducted on the primary inflorescence of *Arabidopsis thaliana* Col-0 growing under continuous light, which are partially reported in Fig. 1 and Supplemental Video 1, are in agreement with the literature. We observed elliptic and circular oscillatory patterns, which occurred in both directions, as well as pendular oscillations. However, the observed inflorescences did not exhibit clear trochoid-like patterns. On the one hand, the existence of pendular circumnutations cannot be explained by the intrinsic oscillator model alone, without *ad hoc* endogenous prescriptions, whereas the flutter instability mechanism might reproduce pendular, elliptic and circular trajectories. On the other hand, a model based on the flutter instability alone seems unable to reproduce the trochoid-like patterns reported in the literature, which would indeed require the superposition of different oscillation modes. Therefore the present study suggests that the preferred hypothesis for the nature of circumnutations should take into account both mechanisms. The relative importance of exogenous versus endogenous oscillations is an emergent property of the system. The first become dominant as the shoot length increases, due to the increasing importance of elastic deformations caused by gravity loading. In other words, the role of elastic deformations in controlling the relative importance of the two mechanisms and the geometry of the oscillations is crucial.

In concluding, we remark that circular trajectories of the plant tip might be the byproduct of having assumed the plant cross section to be circular. Indeed we believe that this is not an intrinsic property of the physical system and preliminary results (not reported) for rods with elliptic cross sections show the emergence of patterns that differ from circular ones. We reserve future studies to explore this observation, together with the need for a quantitative assessment of the accuracy of the theoretical predictions in comparison with experimental observations.

## Data accessibility

A Python implementation of the models is available at https://github.com/mathLab/MorphoelasticRod. This exploits the DOLFIN library as interface for the FEniCS Project Version 2019.1.0 [22], as discussed in the Supplemental Section S5.

## Acknowledgments

The authors would like to thank Alessandro Lucantonio for fruitful discussions and valuable suggestions.

