## Supplementary notes for "Nutations in plant shoots: endogenous and exogenous factors in the presence of mechanical deformations"

#### Contents

**S1 Tip growth** **1**

**S2 Differential growth and evolution laws** **7**

**S3 The statoliths avalanche dynamics** **9**

**S4 Stability analyses** **11**

**S5 Computational model** **19**

**S6 Supplemental Videos** **21**

#### S1 Tip growth

In order to describe the primary growth of plant organs, Erickson and Sax [6] introduced the notion of *relative elemental growth rate* (REGR) or *relative elongation rate* (RER), which in our notation is the material gradient of the Lagrangian velocity field $v_v(S,t) := \partial_t s_v(S,t)$, namely,

$$\mathrm{REGR}(s_v,t) := \mathrm{grad}\, v_v(s_v,t) = \frac{\partial v_v(s_v,t)}{\partial s_v} = \frac{\partial}{\partial s_v}\left(\left.\frac{\partial s_v}{\partial t}\right|_{S(s_v,t)}\right). \tag{S1.1}$$

Such a quantity is related to the deformation gradient, $F = \gamma$, by means of the relationship $\mathrm{grad}\, v_v = \dot{F}F^{-1}$, which explicitly reads

$$\frac{\partial}{\partial s_v}\left(\left.\frac{\partial s_v}{\partial t}\right|_{S(s_v,t)}\right) = \frac{\partial}{\partial t}\left.\left(\frac{\partial s_v}{\partial S}\right)\right|_{S(s_v,t)} \cdot \frac{\partial S}{\partial s_v} = \left.\left(\frac{1}{\gamma}\frac{\partial \gamma}{\partial t}\right)\right|_{S(s_v,t)}, \tag{S1.2}$$

thus yielding

$$\mathrm{REGR}(s_v,t) = \left.\left(\frac{1}{\gamma}\frac{\partial \gamma}{\partial t}\right)\right|_{S(s_v,t)} =: \dot{\varepsilon}_v^\star(s_v,t), \tag{S1.3}$$

where dots denote material time derivatives. Therefore, tip growth is prescribed by the following coupled problems,

$$\frac{\partial s_v}{\partial S}(S,t) = \gamma(S,t) \quad \text{with} \quad s_v(0,t) = 0, \tag{S1.4a}$$

[1] *SISSA–International School for Advanced Studies, 34136 Trieste, Italy*

[2] *The BioRobotics Institute, Scuola Superiore Sant'Anna, 56127 Pisa, Italy*

$$\frac{1}{\gamma(S,t)}\frac{\partial\gamma}{\partial t}(S,t) = \mathrm{REGR}(S,t) \quad \text{with} \quad \gamma(S,0)=1, \tag{S1.4b}$$

for $S \in [0,\ell_0]$ and $t \geq 0$, which can be integrated to get

$$s_v(S,t) = \int_0^S e^{\int_0^t \mathrm{REGR}(\zeta,\tau)\,\mathrm{d}\tau}\,\mathrm{d}\zeta. \tag{S1.5}$$

In addition, if the solution $s_v$ is sufficiently regular, a change of the order of partial derivatives in (S1.4) yields

$$\frac{\partial}{\partial S}\left(\frac{\partial s_v}{\partial t}(S,t)\right) = \mathrm{REGR}(S,t)\frac{\partial s_v}{\partial S}(S,t), \tag{S1.6}$$

so that by integrating first in space and then in time, we arrive at

$$s_v(S,t) = S + \int_0^t \int_0^{s_v(S,\tau)} \mathrm{REGR}(s_v^{-1}(\zeta,\tau),\tau)\,\mathrm{d}\zeta\,\mathrm{d}\tau. \tag{S1.7}$$

As stated in the main text, we assume the relative elongation rate to be a nonnegative function $G$ that vanishes outside the apical growth zone of constant length $\ell_g$, *i.e.*, $G(S,t)=0$ for $S \in [P(t),\ell_0]$ where $P(t) := S(\ell_v(t)-\ell_g,t)$ denotes the material point that exits the growth zone at time $t$. More precisely, we consider

$$\begin{aligned} G(S,t) &= H(S-P(t))F(s_v(S,t)-s_v(P(t),t)) \qquad &\text{(S1.8a)}\\ &= H(s_v(S,t)-(\ell_v(t)-\ell_g))F(s_v(S,t)-(\ell_v(t)-\ell_g)) \qquad &\text{(S1.8b)} \end{aligned}$$

where $H(\cdot)$ is the Heaviside function and $F:[0,\ell_g]\to\mathbb{R}_+$ is a nonzero continuous function. As stated in the following theorem, $P(t)$ is invertible and we denote by $t^\star(S)$ its inverse that is the time at which the material point $S$ exits the growth zone, as its distance from the tip exceeds $\ell_g$.

**Theorem S1.1.** *Let $G$ be a function of the kind* (S1.8). *Then for all $t \geq 0$, $s_v(\cdot,t)$ is monotone increasing (hence invertible). Moreover, $P(t) := s_v^{-1}(s_v(\ell_0,t)-\ell_g,t)$ is monotone increasing (hence invertible).*

*Proof.* In view of equation (S1.5), the function $s_v(\cdot,t)$ is monotone increasing for any fixed time $t$. Moreover, since $G(S,t)\geq 0$, also $\gamma(S,\cdot)$ is an increasing function for any fixed $S\in[0,\ell_0]$. Let us now consider $t_1<t_2$ and the map $f:[0,\ell_v(t_1)]\to[0,\ell_v(t_2)]$ defined as $f(\zeta) := s_v(s_v^{-1}(\zeta,t_1),t_2)$. Since $\gamma(S,\cdot)$ is increasing, we have

$$f'(\zeta) = \frac{\partial s_v}{\partial S}(s_v^{-1}(\zeta,t_1),t_2)\left[\frac{\partial s_v}{\partial S}(s_v^{-1}(\zeta,t_1),t_1)\right]^{-1} = \frac{\gamma(s_v^{-1}(\zeta,t_1),t_2)}{\gamma(s_v^{-1}(\zeta,t_1),t_1)} \geq 1 \tag{S1.9}$$

for all $\zeta\in[0,\ell_v(t_1)]$. Moreover, since $F$ is continuous and nonzero, there exists an interval of positive measure in $[\ell_v(t_1)-\ell_g,\ell_v(t_1)]$ where $f'>1$ so that

$$\ell_v(t_2)-s_v(P(t_2),t_2) = \int_{s_v(P(t_1),t_1)}^{\ell_v(t_1)} \frac{\mathrm{d}f}{\mathrm{d}\zeta}(\zeta)\,\mathrm{d}\zeta > \ell_v(t_1)-s(P(t_1),t_1) = \ell_g \tag{S1.10}$$

that is

$$s_v(P(t_1),t_2) < \ell_0(t_2)-\ell_g = s_v(P(t_2),t_2). \tag{S1.11}$$

Finally, since $s_v(\cdot,t)$ is monotone increasing, we conclude that $P(t_1)<P(t_2)$. □

In addition, for $G$ of the form (S1.8), equation (S1.7) yields

$$\begin{aligned} \ell_v(t) := s_v(\ell_0,t) =& \ell_0 + \int_0^t\int_0^{\ell_v(\tau)} H(\zeta-(\ell_v(\tau)-\ell_g))F(\zeta-\ell_v(\tau)+\ell_g)\,\mathrm{d}\zeta\,\mathrm{d}\tau\\ =& \ell_0 + \int_0^t\int_{\max\{0,\ell_v(\tau)-\ell_g\}}^{\ell_v(\tau)} F(\zeta-\ell_v(\tau)+\ell_g)\,\mathrm{d}\zeta\,\mathrm{d}\tau\\ =& \ell_0 + \int_0^t\int_{\max\{0,\ell_g-\ell_v(\tau)\}}^{\ell_g} F(\zeta)\,\mathrm{d}\zeta\,\mathrm{d}\tau. \end{aligned} \tag{S1.12}$$

so that, for $\ell_0 \geq \ell_g$,

$$\ell_v(t) = \ell_0 + t\int_0^{\ell_g} F(\zeta)\,\mathrm{d}\zeta, \tag{S1.13}$$

which is linear in time, regardless of the function $F$.

In the following we present two cases in which the integral representation (S1.5) can be used to determine an analytical solution to problem (S1.4), which are depicted in Fig. 2a,b of the main text. Then for a few other cases (shown in Fig. 2c,d of the main text and Fig. S1), we solve the problem numerically and we make use of equation (S1.12) to analytically determine $\ell_v(t)$. As for the numerical scheme, we first approximate problem (S1.4b) by

$$\begin{cases} \gamma(S,t_0) = 1, \\ \gamma(S,t_{n+1}) = \gamma(S,t_n)\,[1 + (t_{n+1} - t_n)G(S,t_n)] \quad n \geq 1, \end{cases} \tag{S1.14}$$

where $t_n := nh$ for a sufficiently small time-step $h$, and then we solve for $s_v(S,t_n)$ by integrating $\gamma(S,t_n)$ in space.

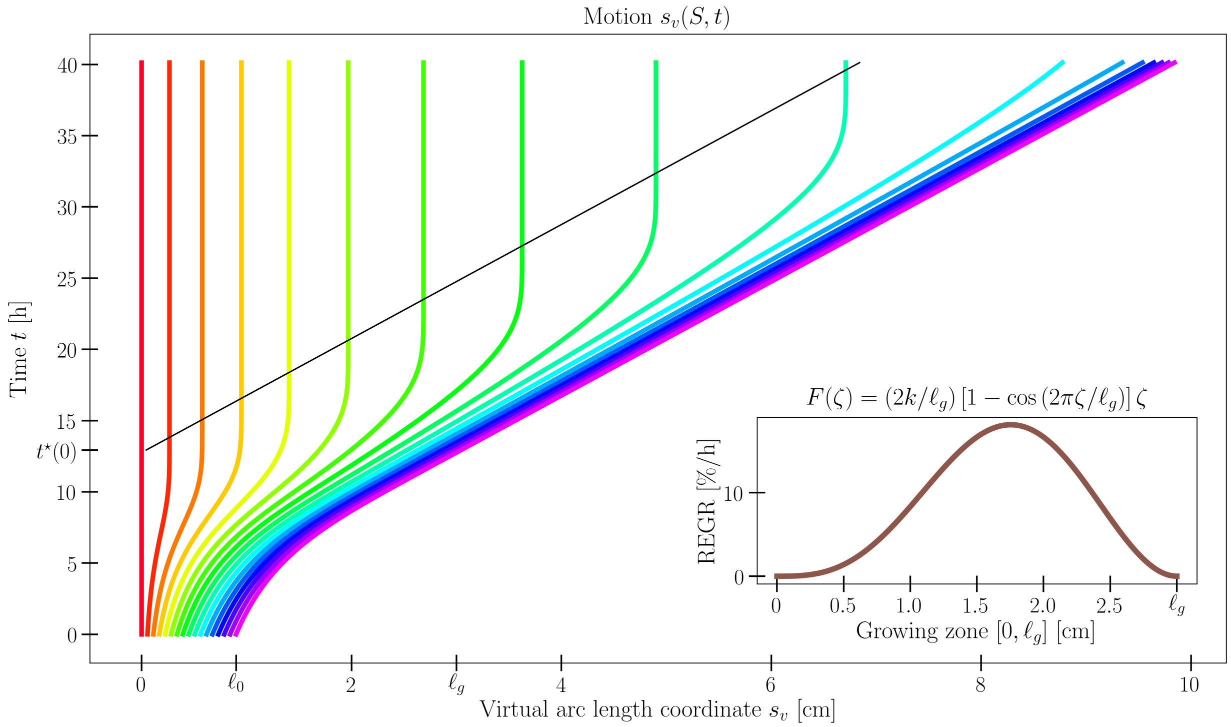

Figure S1: Time evolution of the virtual arc length $s_v(S,t)$ for a set of 17 material points along the plant organ for $G$ as in equation (S1.8) where $F(\zeta) = (2k/\ell_g)\,[1 - \cos(2\pi\zeta/\ell_g)]\,\zeta$. Model parameters are $\ell_0 = 0.9$ cm, $\ell_g = 3$ cm and $k = 0.05$ h$^{-1}$. Notice the black line that denotes the time $t^\star(S)$ at which the material point $S$ exits the growth zone, as its distance from the tip exceeds $\ell_g$. We refer to Supplemental Section S1 for additional details.

**Example 1.**

Consider $G = kH(S - P(t))$ where $k$ is a positive constant. By means of equation (S1.5),

$$\begin{aligned} \ell_v(t) :=& s_v(\ell_0,t) = \int_0^{\ell_0} e^{\int_0^t kH(\zeta - P(\tau))\,\mathrm{d}\tau}\,\mathrm{d}\zeta = \int_0^{P(t)} e^{\int_0^t kH(\zeta - P(\tau))\,\mathrm{d}\tau}\,\mathrm{d}\zeta + \int_{P(t)}^{\ell_0} e^{kt}\,\mathrm{d}\zeta \\ =& s_v(P(t),t) + [\ell_0 - P(t)]\,e^{kt} = \ell_v(t) - \ell_g + [\ell_0 - P(t)]\,e^{kt} \end{aligned} \tag{S1.15}$$

whence

$$P(t) = \ell_0 - \ell_g e^{-kt} \quad \text{and} \quad t^\star(S) = \frac{1}{k}\ln\left(\frac{\ell_g}{\ell_0 - S}\right). \tag{S1.16}$$

We recall that $t^\star(S)$ is the instant of time at which the cell initially located at $S$ stops elongating and we notice that $\lim_{S\to\ell_0} t^\star(S) = \infty$, namely, the tip is never going to stop growing. By combining equations (S1.16) with equation (S1.5), we arrive at

$$s_v(S,t) = [1 - H(S - (\ell_0 - \ell_g))]\,S$$

$$+ H(S - (\ell_0 - \ell_g)) \left\{ H(t - t^\star(S)) \left[\max\{\ell_0, \ell_g\} + \ell_g k \min\{t^\star(S), t^\star(S) - t^\star(0)\} - \ell_g\right] \right.$$
$$+ [1 - H(t - t^\star(S))] \left[ -(\ell_0 - S)e^{kt} + [1 - H(t - t^\star(0))]\ell_0 e^{kt} \right.$$
$$\left.\left. + H(t - t^\star(0)) \left[\max\{\ell_0, \ell_g\} + \ell_g k \min\{t, t - t^\star(0)\}\right] \right] \right\}, \tag{S1.17}$$

whence

$$\ell_v(t) := s_v(\ell_0, t) = \begin{cases} \ell_0 e^{kt} & \text{if } t \le t^\star(0), \\ \max\{\ell_0, \ell_g\} + \ell_g k\,(t - \max\{0, t^\star(0)\}) & \text{if } t > t^\star(0). \end{cases} \tag{S1.18}$$

Finally, equation (S1.17) can be rewritten in the following compact form

$$s_v(S, t) = \begin{cases} S & \text{if } S \le \ell_0 - \ell_g, \\ \ell_v(t^\star(S)) - \ell_g & \text{if } S > \ell_0 - \ell_g \text{ and } t \ge t^\star(S), \\ \ell_v(t) - (\ell_0 - S)e^{kt} & \text{if } S > \ell_0 - \ell_g \text{ and } t < t^\star(S), \end{cases} \tag{S1.19}$$

which is shown in Fig. 2a of the main text.

**Example 2.**

Drawing inspiration from the REGR profiles experimentally measured in growing roots [4, 16], we consider $G$ as in equation (S1.8) with $F(\zeta) = k + (k_1 - k)H(\zeta - (\ell_g - \ell_1))$ where $k$, $k_1 > 0$ and $0 < \ell_1 < \ell_g$. Upon defining

$$P_1(t) := s_v^{-1}(s_v(\ell_0, t) - \ell_1, t), \tag{S1.20}$$

we get

$$G(S,t) = H(S - P(t))\left[k_1 H(S - P_1(t)) + k\,(1 - H(S - P_1(t)))\right] = \begin{cases} 0 & \text{if } S \le P(t), \\ k & \text{if } P(t) < S \le P_1(t), \\ k_1 & \text{if } P_1(t) < S \le \ell_v(t). \end{cases} \tag{S1.21}$$

We first determine the functions $P$ and $P_1$ together with their inverse. We notice that

$$\begin{aligned} \ell_v(t) :=& s_v(\ell_0, t) = \int_0^S e^{\int_0^t H(\zeta - P(\tau))[k_1 H(\zeta - P_1(\tau)) + k(1 - H(\zeta - P_1(\tau)))]\,\mathrm{d}\tau}\,\mathrm{d}\zeta \\ =& s_v(P_1(t), t) + \int_{P_1(t)}^S e^{k_1 t}\,\mathrm{d}\zeta = \ell_v(t) - \ell_1 + \left[\ell_0 - P_1(t)\right] e^{k_1 t}, \end{aligned} \tag{S1.22}$$

whence

$$P_1(t) = \ell_0 - \ell_1 e^{-k_1 t} \quad \text{and} \quad t_1^\star(S) = \frac{1}{k_1}\ln\left(\frac{\ell_1}{\ell_0 - S}\right). \tag{S1.23}$$

Moreover,

$$\begin{aligned} \ell_v(t) :=& s_v(\ell_0, t) = \int_0^S e^{\int_0^t H(\zeta - P(\tau))[k_1 H(\zeta - P(\tau)) + k(1 - H(\zeta - P_1(\tau)))]\,\mathrm{d}\tau}\,\mathrm{d}\zeta \\ =& s_v(P(t), t) + H(P(t) - (\ell_0 - \ell_1)) \int_{P(t)}^{P_1(t)} e^{k_1 t + k(t - t_1^\star(\zeta))}\,\mathrm{d}\zeta \\ & + [1 - H(P(t) - (\ell_0 - \ell_1))] \left[ \int_{P(t)}^{\ell_0 - \ell_1} e^{kt}\,\mathrm{d}\zeta + \int_{\ell_0 - \ell_1}^{P_1(t)} e^{k_1 t_1^\star(\zeta) + k(t - t_1^\star(\zeta))}\,\mathrm{d}\zeta \right] + \int_{P_1(t)}^{\ell_0} e^{k_1 t}\,\mathrm{d}\zeta \\ =& \ell_v(t) - \ell_g + H(P(t) - (\ell_0 - \ell_1)) \int_{P(t)}^{P_1(t)} e^{kt} \left(\frac{\ell_1}{\ell_0 - \zeta}\right)^{1 - \frac{k}{k_1}} \mathrm{d}\zeta \\ & + [1 - H(P(t) - (\ell_0 - \ell_1))] \left[ (\ell_0 - \ell_1 - P(t))e^{kt} + \int_{\ell_0 - \ell_1}^{P_1(t)} e^{kt} \left(\frac{\ell_1}{\ell_0 - \zeta}\right)^{1 - \frac{k}{k_1}} \mathrm{d}\zeta \right] + \left[\ell_0 - P_1(t)\right] e^{k_1 t} \\ =& \ell_v(t) - (\ell_g - \ell_1) + H(P(t) - (\ell_0 - \ell_1))\ell_1 \frac{k_1}{k} \left[ \left(\frac{\ell_0 - P(t)}{\ell_1}\right)^{k/k_1} e^{kt} - 1 \right] \\ & + [1 - H(P(t) - (\ell_0 - \ell_1))] \left[ (\ell_0 - \ell_1 - P(t))e^{kt} + \ell_1 \frac{k}{k_1}\left(e^{kt} - 1\right) \right] \end{aligned}$$

$$=\ell_v(t)+\begin{cases}\ell_1\left[1-\frac{k_1}{k}-\left(1-\frac{k_1}{k}\right)e^{kt}\right]+(\ell_0-P(t))e^{kt}-\ell_g & \text{if } P(t)<\ell_0-\ell_1,\\ \ell_1\left[1-\frac{k_1}{k}+\frac{k_1}{k}\left(\frac{\ell_0-P}{\ell_1}\right)^{\frac{k}{k_1}}e^{kt}\right]-\ell_g & \text{if } P(t)\geq\ell_0-\ell_1.\end{cases} \tag{S1.24}$$

Then we deduce that

$$P(t)=\begin{cases}\ell_0-\left[\ell_g-\ell_1\left(1-\frac{k_1}{k}(1-e^{kt})\right)\right]e^{-kt} & \text{if } t\leq\frac{1}{k}\ln\left(1+\frac{k}{k_1}\left(\frac{\ell_g}{\ell_1}-1\right)\right),\\ \ell_0-\ell_1\left[1+\frac{k}{k_1}\left(\frac{\ell_g}{\ell_1}-1\right)\right]^{\frac{k_1}{k}}e^{-k_1t} & \text{otherwise},\end{cases} \tag{S1.25}$$

whose inverse is

$$t^\star(S)=\begin{cases}\frac{1}{k}\ln\left(\frac{k_1\ell_1+k(\ell_g-\ell_1)}{k_1\ell_1+k(\ell_0-\ell_1-S)}\right) & \text{if } S\leq\ell_0-\ell_1,\\ \frac{1}{k}\ln\left(1+\frac{k}{k_1\ell_1}(\ell_g-\ell_1)\right)+\frac{1}{k_1}\ln\left(\frac{\ell_1}{\ell_0-S}\right) & \text{if } S>\ell_0-\ell_1.\end{cases} \tag{S1.26}$$

We next notice that

$$\begin{aligned}\int_0^t G(\zeta,\tau)\,\mathrm{d}\tau &= \int_0^t H(\zeta-P(\tau))\left[k_1H(\zeta-P_1(\tau))+k\left(1-H(\zeta-P_1(\tau))\right)\right]\mathrm{d}\tau\\ &=\begin{cases}0 & \text{if } \zeta\leq\ell_0-\ell_g,\\ kt^\star(\zeta) & \text{if } \ell_0-\ell_g<\zeta\leq\min\{\ell_0-\ell_1,P(t)\},\\ kt & \text{if } P(t)<\zeta\leq\ell_0-\ell_1,\\ k_1t_1^\star(\zeta)+k[t^\star(\zeta)-t_1^\star(\zeta)] & \text{if } \ell_0-\ell_1<\zeta\leq P(t),\\ k_1t_1^\star(\zeta)+k[t-t_1^\star(\zeta)] & \text{if } \max\{P(t),\ell_0-\ell_1\}<\zeta\leq P_1(t),\\ k_1t & \text{if } \zeta>P_1(t),\end{cases}\end{aligned} \tag{S1.27}$$

and equation (S1.5) can be used to deduce the following expression for the total virtual length

$$\ell_v(t)=\begin{cases}\ell_0e^{k_1t} & t\leq t_1^\star(0),\\ \left[\max\{\ell_0-\ell_1,0\}+\ell_1\frac{k_1}{k}\right]e^{k(t-\max\{0,t_1^\star(0)\})}+\ell_1(1-\frac{k_1}{k}) & t_1^\star(0)<t\leq t^\star(0),\\ \max\{\ell_0,\ell_g\}+\left[(\ell_g-\ell_1)k+\ell_1k_1\right](t-\max\{0,t^\star(0)\}) & t>t^\star(0),\end{cases} \tag{S1.28}$$

for any $t>0$. Finally, we arrive at

$$s_v(S,t)=\begin{cases}S & \text{if } S\leq\ell_0-\ell_g,\\ \ell_v(t)-(\ell_0-S)e^{k_1t} & \text{if } S>\ell_0-\ell_g \text{ and } t\leq t_1^\star(S),\\ \ell_v(t)-\ell_g+(S-P(t))e^{kt} & \text{if } S>\ell_0-\ell_g \text{ and } t_1^\star(S)<t\leq t^\star(S)\leq t^\star(\ell_0-\ell_1),\\ \ell_v(t)-\ell_1-\ell_1\frac{k_1}{k}\left[e^{k(t-t_1^\star(S))}-1\right] & \text{if } S>\ell_0-\ell_g,\ t_1^\star(S)<t\leq t^\star(S), \text{ and } t^\star(S)>t^\star(\ell_0-\ell_1)\\ \ell_v(t^\star(S))-\ell_g & \text{if } S>\ell_0-\ell_g \text{ and } t>t^\star(S),\end{cases} \tag{S1.29}$$

which is illustrated in Fig. 2b of the main text.

**Example 3.**

Inspired by the REGR profiles experimentally measured in growing *Arabidopsis thaliana* inflorescence stems by Phyo et al. [13], we consider $G$ as in equation (S1.8) with a linear function $F(\zeta)=2k\zeta/\ell_g$ where $k>0$. For $\ell_g\leq\ell_0$, we have $\ell_g<\ell_v(t)$ so that

$$\ell_v(t)=\ell_0+\int_0^t\int_0^{\ell_g}F(\zeta)\,\mathrm{d}\zeta\,\mathrm{d}\tau=\ell_0+\frac{k}{\ell_g}\int_0^t\int_0^{\ell_g}2\zeta\,\mathrm{d}\zeta\,\mathrm{d}\tau=\ell_0+\ell_gkt. \tag{S1.30}$$

Fig. 2c in the main text shows the numerical solution of the problem (S1.4) corresponding to such a choice.

**Example 4.**

Drawing inspiration from the REGR profiles measured by Hall and Ellis [9], we consider $G$ as in equation (S1.8) with $F(\zeta)=k\left(1-\cos\left(2\pi\zeta/\ell_g\right)\right)$ where $k>0$. By assuming that $\ell_g\leq\ell_0$, we get

$$\begin{aligned}\ell_v(t)=&\ell_0+\int_0^t\int_0^{\ell_g}F(\zeta)\,\mathrm{d}\zeta\,\mathrm{d}\tau=\ell_0+kt\int_0^{\ell_g}\left[1-\cos\left(\frac{2\pi\zeta}{\ell_g}\right)\right]\mathrm{d}\zeta\\ =&\ell_0+kt\left[\zeta-\frac{\ell_g}{2\pi}\sin\left(\frac{2\pi\zeta}{\ell_g}\right)\right]_0^{\ell_g}\mathrm{d}\zeta=\ell_0+\ell_gkt.\end{aligned} \tag{S1.31}$$

A numerical solution of the problem (S1.4) corresponding to such a choice is illustrated in Fig. 2d of the main text.

**Example 5.**

Consider $G$ as in equation (S1.8) with $F(\zeta) = (2k/\ell_g)\,[1 - \cos(2\pi\zeta/\ell_g)]\,\zeta$ where $k > 0$. For $\ell_g \leq \ell_0$, we get

$$\begin{aligned}\ell_v(t) =& \ell_0 + \int_0^t \int_0^{\ell_g} F(\zeta)\,\mathrm{d}\zeta\,\mathrm{d}\tau = \ell_0 + \frac{k}{\ell_g} t \int_0^{\ell_g} 2\zeta \left[1 - \cos\left(\frac{2\pi\zeta}{\ell_g}\right)\right] \mathrm{d}\zeta \\ =& \ell_0 + \frac{k}{\ell_g} t \left[\zeta^2 - \frac{\ell_g^2}{2\pi^2} \cos\left(\frac{2\pi\zeta}{\ell_g}\right) - \frac{\zeta\ell_g}{\pi} \sin\left(\frac{2\pi\zeta}{\ell_g}\right)\right]_0^{\ell_g} \\ =& \ell_0 + \ell_g k t. \end{aligned} \tag{S1.32}$$

Fig. S1 shows $s_v(S,t)$ as numerically computed by means of the numerical scheme (S1.14).

#### S2 Differential growth and evolution laws

In this section we derive expressions (2.13)-(2.14) of the main text, which determine the relationship between differential growth and strain rates. We start by extending the notion of relative elemental growth rate to any point laying on the circular cross section $s_v$ of the virtual configuration at time $t$. We parameterize the surface by means of the spatial coordinates $(x, y)$ in the local basis $\{\mathbf{d}_1^v(s_v, t), \mathbf{d}_2^v(s_v, t)\}$, namely,

$$\mathbf{p}_v(s_v, t; x, y) := \mathbf{p}_v(s_v, t) + x\,\mathbf{d}_1^v(s_v, t) + y\,\mathbf{d}_2^v(s_v, t). \tag{S2.1}$$

Then the length of the material fiber passing through point $(x, y)$ of the cross section $s_v$ and extending from the rod's base to that point, can be written as

$$\begin{aligned} \ell_v(s_v, t; x, y) &:= \int_0^{s_v} \left(\partial_{s_v}\mathbf{p}_v(\zeta, t; x, y) \cdot \partial_{s_v}\mathbf{p}_v(\zeta, t; x, y)\right)^{\frac{1}{2}} \mathrm{d}\zeta \\ &= \int_0^{s_v} \left\{[1 + u_1^\star y - u_2^\star x]^2 + (x^2 + y^2)\, {u_3^\star}^2\right\}^{\frac{1}{2}} \mathrm{d}\zeta. \end{aligned} \tag{S2.2}$$

Equation (S2.2) follows by using the kinematic relationships $\partial_{s_v}\mathbf{p}_v = \mathbf{d}_3^v$ and $\partial_{s_v}\mathbf{d}_i^v = \mathbf{u}^\star \times \mathbf{d}_i^v\ \forall\, i$, where $\mathbf{u}^\star = \sum_j u_j^\star \mathbf{d}_j^v$ is the spontaneous twist. Then the growth stretch at $(x, y)$ is given by

$$\begin{aligned} \gamma(s_v, t; x, y) :=& \left.\frac{\partial \ell_v(s_v, t; x, y)}{\partial S}\right|_{S=S(s_v,t)} = \left.\frac{\partial \ell_v(s_v, t; x, y)}{\partial s_v}\frac{\partial s_v}{\partial S}\right|_{S=S(s_v,t)} \\ =& \gamma(s_v, t) \left.\left\{[1 + u_1^\star y - u_2^\star x]^2 + \left(x^2 + y^2\right) {u_3^\star}^2\right\}^{\frac{1}{2}}\right|_{(s_v,t)}, \end{aligned} \tag{S2.3}$$

so that the true strain rate reads

$$\dot{\varepsilon}_v^\star(s_v, t; x, y) = \frac{\dot{\gamma}}{\gamma}(s_v, t; x, y) = \dot{\varepsilon}_v^\star(s_v, t) + \frac{\left[(1 + u_1^\star y - u_2^\star x)\left(\dot{u}_1^\star y - \dot{u}_2^\star x\right) + u_3^\star \dot{u}_3^\star \left(x^2 + y^2\right)\right]}{\left(1 + u_1^\star y - u_2^\star x\right)^2 + {u_3^\star}^2\left(x^2 + y^2\right)}. \tag{S2.4}$$

By differentiating expression (S2.4) with respect to $x$ and $y$, we get

$$\begin{aligned} \partial_x \dot{\varepsilon}_v^\star(s_v, t; x, y) =& \frac{\left[-\dot{u}_2^\star - \left(u_1^\star \dot{u}_2^\star + u_2^\star \dot{u}_1^\star\right) y + 2\left(u_2^\star \dot{u}_2^\star + u_3^\star \dot{u}_3^\star\right) x\right]\left[\left(1 + u_1^\star y - u_2^\star x\right)^2 + {u_3^\star}^2\left(x^2 + y^2\right)\right]}{\left[\left(1 + u_1^\star y - u_2^\star x\right)^2 + {u_3^\star}^2\left(x^2 + y^2\right)\right]^2} \\ &- \frac{\left[(1 + u_1^\star y - u_2^\star x)\left(\dot{u}_1^\star y - \dot{u}_2^\star x\right) + u_3^\star \dot{u}_3^\star\left(x^2 + y^2\right)\right]\left[-2u_2^\star\left(1 + u_1^\star y - u_2^\star x\right) + 2{u_3^\star}^2 x\right]}{\left[\left(1 + u_1^\star y - u_2^\star x\right)^2 + {u_3^\star}^2\left(x^2 + y^2\right)\right]^2}, \end{aligned} \tag{S2.5}$$

and

$$\begin{aligned} \partial_y \dot{\varepsilon}_v^\star(s_v, t; x, y) =& \frac{\left[\dot{u}_1^\star - \left(u_1^\star \dot{u}_2^\star + u_2^\star \dot{u}_1^\star\right) x + 2\left(u_1^\star \dot{u}_1^\star + u_3^\star \dot{u}_3^\star\right) y\right]\left[\left(1 + u_1^\star y - u_2^\star x\right)^2 + {u_3^\star}^2\left(x^2 + y^2\right)\right]}{\left[\left(1 + u_1^\star y - u_2^\star x\right)^2 + {u_3^\star}^2\left(x^2 + y^2\right)\right]^2} \\ &- \frac{\left[(1 + u_1^\star y - u_2^\star x)\left(\dot{u}_1^\star y - \dot{u}_2^\star x\right) + u_3^\star \dot{u}_3^\star\left(x^2 + y^2\right)\right]\left[2u_1^\star\left(1 + u_1^\star y - u_2^\star x\right) + 2{u_3^\star}^2 y\right]}{\left[\left(1 + u_1^\star y - u_2^\star x\right)^2 + {u_3^\star}^2\left(x^2 + y^2\right)\right]^2}, \end{aligned} \tag{S2.6}$$

respectively. Therefore, by Taylor expanding (S2.4) about the cross section center $(0, 0)$, we arrive at

$$\begin{aligned} \dot{\varepsilon}_v^\star(s_v, t; x, y) &= \dot{\varepsilon}_v^\star(s_v, t) + \nabla\dot{\varepsilon}_v^\star(s_v, t; 0, 0) \cdot \left(x\,\mathbf{d}_1^v(s_v, t) + y\,\mathbf{d}_2^v(s_v, t)\right) + o(\sqrt{x^2 + y^2}) \\ &= \dot{\varepsilon}_v^\star(s_v, t) + \left[-\dot{u}_2^\star(s_v, t), \dot{u}_1^\star(s_v, t)\right] \cdot [x, y] + o(\sqrt{x^2 + y^2}), \end{aligned} \tag{S2.7}$$

from which equations (2.13)-(2.14) in the main text follow.

**Remark S2.1.** Prescribing the *growth gradient* $\boldsymbol{\delta}_v := \nabla\dot{\varepsilon}_v^\star(s_v, t; 0, 0)$ is equivalent to the approaches taken in previous studies [2, 3, 14], which involve a notion of *differential growth* $\mathrm{DG}(s_v, t; \vartheta)$ introduced as a means to compare strains at diametrically opposite sides of the circular cross section. Indeed, by passing to the polar coordinates $(\rho, \vartheta)$, such that $(x, y) = (\rho\cos\vartheta, \rho\sin\vartheta)$, equation (S2.4) reads

$$\dot{\varepsilon}_v^\star(s_v, t; \rho, \vartheta) = \dot{\varepsilon}_v^\star(s_v, t) + \frac{\left[(1 + u_1^\star\rho\sin\vartheta - u_2^\star\rho\cos\vartheta)\,(\dot{u}_1^\star\rho\sin\vartheta - \dot{u}_2^\star\rho\cos\vartheta) + u_3^\star\dot{u}_3^\star\rho^2\right]}{(1 + u_1^\star\rho\sin\vartheta - u_2^\star\rho\cos\vartheta)^2 + {u_3^\star}^2\rho^2}, \tag{S2.8}$$

and then, the differential growth can be defined as

$$\mathrm{DG}(s_v, t; \vartheta) := \frac{\dot{\varepsilon}_v^\star(s_v, t; r, \vartheta) - \dot{\varepsilon}_v^\star(s_v, t; r, \vartheta + \pi)}{\dot{\varepsilon}_v^\star(s_v, t; r, \vartheta) + \dot{\varepsilon}_v^\star(s_v, t; r, \vartheta + \pi)} = \frac{(\dot{a} - \dot{b})\,(A + (b - a)B) - c\dot{c}B}{\dot{\varepsilon}_v^\star(s_v, t)\,(A^2 - B^2) + (\dot{a} - \dot{b})\,((a - b)A - B) + c\dot{c}}, \tag{S2.9}$$

or

$$\mathrm{DG}(s_v, t; \vartheta) := \frac{\dot{\varepsilon}_v^\star(s_v, t; r, \vartheta) - \dot{\varepsilon}_v^\star(s_v, t; r, \vartheta + \pi)}{2\dot{\varepsilon}_v^\star(s_v, t)} = \frac{(\dot{a} - \dot{b})\,(A + (b - a)B) - c\dot{c}B}{\dot{\varepsilon}_v^\star(s_v, t)\,(A^2 - B^2)}, \tag{S2.10}$$

where $a := u_1^\star r\sin\vartheta$, $b := u_2^\star r\cos\vartheta$, $c := u_3^\star r$, $A := 1 + (a - b)^2 + c^2$ and $B := 2(a - b)$. In both cases, by assuming that $ru_j^\star \ll 1$, the differential growth DG can be approximated as

$$\mathrm{DG}(s_v, t; \vartheta) \simeq \frac{r}{\dot{\varepsilon}_v^\star}(\dot{u}_1^\star\sin\vartheta - \dot{u}_2^\star\cos\vartheta) = \frac{r}{\dot{\varepsilon}_v^\star}\,(\dot{u}_1^\star\mathbf{d}_2^v - \dot{u}_2^\star\mathbf{d}_1^v)\cdot\mathbf{a}(s_v, t; \vartheta), \tag{S2.11}$$

where $\mathbf{a}(s_v, t; \vartheta) := \cos\vartheta\,\mathbf{d}_1^v(s_v, t) + \sin\vartheta\,\mathbf{d}_2^v(s_v, t)$. Therefore, by comparing expressions (S2.7) and (S2.11), we deduce that prescribing the *differential growth* $\mathrm{DG}(s_v, t; \vartheta)$ for all $\vartheta \in [0, 2\pi)$ is equivalent to prescribe the *growth gradient* $\boldsymbol{\delta}_v := \nabla\dot{\varepsilon}_v^\star(s_v, t; 0, 0)$, as introduced in the main text by means of equation (2.14).

#### S3 The statoliths avalanche dynamics

If compared to the purely phenomenological model of Sachs' sine law [17], plant gravitropic responses can be refined by including the dynamics of the statoliths avalanche in plant cells. This has been demonstrated to be the microscopic mechanism through which plant shoots and roots perceive the direction of gravity [11, 5]. We assume the free surface of the statoliths pile in a statocyte cell to be planar and the normal given by the unit vector $\mathbf{h}(s,t)$, as depicted in Fig. S2. Since our reasoning holds for any fixed cross section coordinate $s$, in the following we omit the dependence on such a parameter.

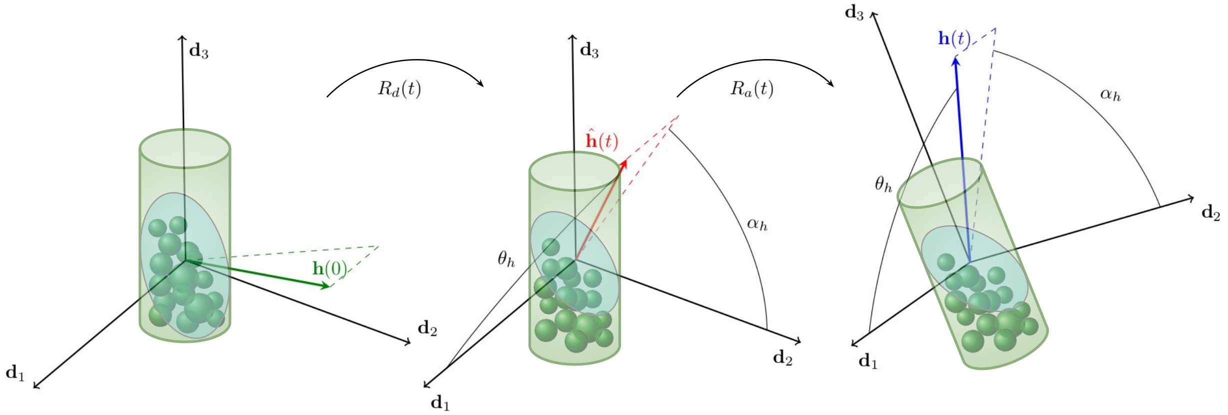

Figure S2: Statoliths avalanche dynamics in a statocyte. The motion of the free surface of piled statoliths can be decomposed into two rotations as in (S3.1): $\hat{\mathbf{h}}(t) = \mathbf{R}_a(t)\mathbf{h}(0)$ is the orientation as described by an observer co-moving with the directors and $\mathbf{h}(t) = \mathbf{R}_d(t)\hat{\mathbf{h}}(t)$ is the orientation as seen by an external observer. Here the unit vector $\mathbf{h}$ is parameterized by two angles defined with respect to the directors: $\theta_h$ is the angle between $\mathbf{h}$ and $\mathbf{d}_1$ and $\alpha_h$ is the angle between $(\mathbf{I} - \mathbf{d}_1 \otimes \mathbf{d}_1)\mathbf{h}$ and $\mathbf{d}_2$.

Fig. S2 shows the decomposition of the motion of $\mathbf{h}$ into two dynamics, namely,

$$\mathbf{h}(t) = \mathbf{R}_d(t)\mathbf{R}_a(t)\mathbf{h}(0), \tag{S3.1}$$

where, for any $t$, $\mathbf{R}_a(t)$ and $\mathbf{R}_d(t)$ are rotations. Here, $\mathbf{R}_a(t)$ can be thought of as the viscous relaxation of the statoliths pile relative to the statocyte so that $\hat{\mathbf{h}}(t) := \mathbf{R}_a(t)\mathbf{h}(0)$ is the dynamics as described by an observer co-moving with the directors, while $\mathbf{R}_d(t)$ is the rotation of the material frame, that is, of the statocyte itself. We prescribe the dynamics of $\hat{\mathbf{h}}$ as a viscous relaxation towards $\mathbf{R}_d(t)^T\mathbf{e}_2$, *i.e.*,

$$\dot{\hat{\mathbf{h}}}(t) = -\frac{1}{\tau_a}\left(\mathbf{R}_d(t)^T\mathbf{e}_2 \times \hat{\mathbf{h}}(t)\right) \times \hat{\mathbf{h}}(t), \tag{S3.2}$$

where $\tau_a$ is the characteristic time scale for the statoliths avalanche dynamics. By taking the time derivative of equation (S3.1) and making use of equation (S3.2), we arrive at

$$\begin{aligned}
\dot{\mathbf{h}}(t) &= \left(\dot{\mathbf{R}}_d(t)\mathbf{R}_a(t) + \mathbf{R}_d(t)\dot{\mathbf{R}}_a(t)\right)\mathbf{h}(0) \\
&= \dot{\mathbf{R}}_d(t)\mathbf{R}_d(t)^T\mathbf{h}(t) + \mathbf{R}_d(t)\dot{\mathbf{R}}_a(t)\mathbf{R}_a(t)^T\hat{\mathbf{h}}(t) \\
&= \mathbf{w}(t) \times \mathbf{h}(t) + \mathbf{R}_d(t)\left[-\frac{1}{\tau_a}\left(\mathbf{R}_d(t)^T\mathbf{e}_2 \times \hat{\mathbf{h}}(t)\right) \times \mathbf{R}_d(t)^T\mathbf{h}(t)\right] \\
&= \left(\mathbf{w}(t) + \frac{1}{\tau_a}\mathbf{h}(t) \times \mathbf{e}_2\right) \times \mathbf{h}(t),
\end{aligned} \tag{S3.3}$$

where $\mathbf{w}(t)$ is the spin, namely, the axial vector associated with $\mathbf{R}_d(t)$. In other terms,

$$\dot{\mathbf{h}}(t) - \mathbf{w}(t) \times \mathbf{h}(t) = \frac{1}{\tau_a}(\mathbf{h}(t) \times \mathbf{e}_2) \times \mathbf{h}(t), \tag{S3.4}$$

and, by using the kinematic relationship $\dot{\mathbf{d}}_j = \mathbf{w} \times \mathbf{d}_j$, we get

$$\sum_j \dot{h}_j(t)\mathbf{d}_j(t) = \frac{1}{\tau_a}(\mathbf{h}(t) \times \mathbf{e}_2) \times \mathbf{h}(t), \tag{S3.5}$$

where $h_j := \mathbf{h} \cdot \mathbf{d}_j$. In terms of components, equation (S3.5) reads

$$\begin{aligned} \dot{h}_j(t) &= \frac{1}{\tau_a} \left[ d_{j2}(t) - (\mathbf{h}(t) \cdot \mathbf{e}_2)(\mathbf{h}(t) \cdot \mathbf{d}_j(t)) \right] \\ &= \frac{1}{\tau_a} \left[ d_{j2}(t) - h_j(t) \sum_i h_i(t) d_{i2}(t) \right] \qquad \forall j, \end{aligned} \tag{S3.6}$$

where $d_{ij} := \mathbf{d}_i \cdot \mathbf{e}_j$ for all $i$, $j = 1, 2, 3$. However, we notice that the components of $\mathbf{h}$ are not independent one from the other, due to the constraint on the norm. *i.e.*, $||\mathbf{h}|| = 1$. Then, from the practical point of view, it is convenient to parameterize $\mathbf{h}$ with two angles defined with respect to a certain frame of reference. One possibility is to consider the angles that $\mathbf{h}$ forms with $\mathbf{d}_1$ and $\mathbf{d}_2$, as shown in Fig. S2. In this case, $\theta_h$ is the angle between $\mathbf{h}$ and $\mathbf{d}_1$, while $\alpha_h$ is the angle between $(\mathbf{I} - \mathbf{d}_1 \otimes \mathbf{d}_1)\mathbf{h}$ and $\mathbf{d}_2$, so that

$$\mathbf{h} = \cos\theta_h \mathbf{d}_1 + \sin\theta_h \cos\alpha_h \mathbf{d}_2 + \sin\theta_h \sin\alpha_h \mathbf{d}_3. \tag{S3.7}$$

Then

$$\dot{h}_1 = -\dot{\theta}_h \sin\theta_h, \tag{S3.8a}$$

$$\dot{h}_2 = \dot{\theta}_h \cos\theta_h \cos\alpha_h - \dot{\alpha}_h \sin\theta_h \sin\alpha_h, \tag{S3.8b}$$

$$\dot{h}_3 = \dot{\theta}_h \cos\theta_h \sin\alpha_h + \dot{\alpha}_h \sin\theta_h \cos\alpha_h, \tag{S3.8c}$$

whence

$$\cos\theta_h \left( \cos\alpha_h \dot{h}_2 + \sin\alpha_h \dot{h}_3 \right) - \sin\theta_h \dot{h}_1 = \dot{\theta}_h, \tag{S3.9a}$$

$$\cos\alpha_h \dot{h}_3 - \sin\alpha_h \dot{h}_2 = \dot{\alpha}_h \sin\theta_h. \tag{S3.9b}$$

Therefore, for $\sin\theta_h \neq 0$, equations (S3.9) provide the evolution laws (S3.5) in terms of the angles $\theta_h$ and $\alpha_h$.

### S4 Stability analyses

In this section we report the stability analyses of the model discussed in Section 3 of the main text. We start by summarizing the governing equations for the model of growing plant shoots. Then, upon introducing their representation in terms of Euler angles, we proceed to study the reduced model described in Section 3.1 of the main text, by gradually exploring the effects of gravitropic and proprioceptive responses.

#### S4.1 Summary of the model for growing plant shoots

Under assumptions (i)-(vii) discussed in Section 2.6 of the main text, we derive a model suitable for the study of growing plant shoots. As for the governing equations, these read

$$\frac{\partial s}{\partial S}(S,t) = \lambda(S,t), \tag{S4.1a}$$

$$\frac{1}{\lambda(S,t)}\frac{\partial \lambda}{\partial t}(S,t) = \begin{cases} 0 & \text{if } s(S,t) \leq \ell(t) - \ell_g, \\ \frac{1}{\tau_g} & \text{if } s(S,t) > \ell(t) - \ell_g, \end{cases} \tag{S4.1b}$$

$$\mathbf{m}'(s,t) = -\,q\,(\ell(t) - s)\,\mathbf{e}_2 \times \mathbf{d}_3(s,t), \tag{S4.1c}$$

$$\mathbf{m} = \sum_j K_j \left(u_j - u_j^\star\right)\mathbf{d}_j, \tag{S4.1d}$$

$$E(s,t) = E_1 - (E_1 - E_0)\, e^{-\frac{1}{\tau_\ell}\max\{0, t - t^\star(S(s,t))\}}, \tag{S4.1e}$$

$$\sum_j \dot{h}_j(s,t)\mathbf{d}_j(s,t) = \frac{1}{\tau_a}(\mathbf{h}(s,t)\times\mathbf{e}_2)\times\mathbf{h}(s,t), \tag{S4.1f}$$

$$\begin{aligned}\dot{u}_1^\star(s,t) =& \alpha\frac{\dot{\varepsilon}^\star(s,t)}{r}\cos(2\pi t/\tau_e) - \beta\frac{\dot{\varepsilon}^\star(s,t)}{r\tau_m}\int_{-\infty}^{t-\tau_r} e^{-\frac{1}{\tau_m}(t-\tau_r-\tau)} h_2(s,\tau)\,\mathrm{d}\tau \\ &- \eta\frac{\dot{\varepsilon}^\star(s,t)}{\bar{\tau}_m}\int_{-\infty}^{t-\bar{\tau}_r} e^{-\frac{1}{\bar{\tau}_m}(t-\bar{\tau}_r-\tau)} u_1(s,\tau)\,\mathrm{d}\tau,\end{aligned} \tag{S4.1g}$$

$$\begin{aligned}\dot{u}_2^\star(s,t) =& \alpha\frac{\dot{\varepsilon}^\star(s,t)}{r}\sin(2\pi t/\tau_e) + \beta\frac{\dot{\varepsilon}^\star(s,t)}{r\tau_m}\int_{-\infty}^{t-\tau_r} e^{-\frac{1}{\tau_m}(t-\tau_r-\tau)} h_1(s,\tau)\,\mathrm{d}\tau \\ &- \eta\frac{\dot{\varepsilon}^\star(s,t)}{\bar{\tau}_m}\int_{-\infty}^{t-\bar{\tau}_r} e^{-\frac{1}{\bar{\tau}_m}(t-\bar{\tau}_r-\tau)} u_2(s,\tau)\,\mathrm{d}\tau,\end{aligned} \tag{S4.1h}$$

where all variables and parameters are defined as in the main text, primes denote differentiation with respect to the parameter $s$ and dots denote material time derivatives. We recall that equations (S4.1a)-(S4.1b) define the tip growth law, equation (S4.1c) follows from the balance of linear and angular momentum where $\mathbf{m}$ is the resultant contact couple given by the constitutive law of (S4.1d). Equation (S4.1e) is the lignification law, equation (S4.1f) governs the statoliths avalanche dynamics, and equations (S4.1g)-(S4.1h) are the evolution laws for the spontaneous strains.

#### S4.2 Representation in terms of Euler angles

The nine components of the directors $\{\mathbf{d}_j\}$ are not independent, due to the orthonormality constraints. Then it is possible to represent the directors in terms of three independent angles, the Euler angles, so that the orthonormality constraints are automatically fulfilled. Although this representation introduces a polar singularity leading to an ambiguity of the representation, this can be successfully adopted in our setting, upon a careful choice of the notation for the Euler angles. We describe the rotation mapping the fixed basis $\{\mathbf{e}_j\}$ to the basis of directors $\{\mathbf{d}_j\}$ by means of the following three successive rotations:

(i) A rotation by an angle $\varphi$ about the $\mathbf{e}_3$-axis;
(ii) A rotation by an angle $\psi$ about the rotated $\mathbf{e}_2$-axis, denoted by $\mathbf{e}_2'$;
(iii) A rotation by an angle $\chi$ about the rotated $\mathbf{e}_3$-axis, denoted by $\mathbf{e}_3''$.

Such a decomposition is illustrated in Fig. S3 and it is well defined if $\psi \neq 0$, otherwise the Euler angles are not uniquely determined by the directors, since only the sum $\chi + \varphi$ can be established. On the other hand, the directors are always uniquely determined by the three angles as

$$\mathbf{d}_1 = (\cos\chi\cos\psi\cos\varphi - \sin\chi\sin\varphi)\,\mathbf{e}_1 + (\cos\chi\cos\psi\sin\varphi + \sin\chi\cos\varphi)\,\mathbf{e}_2 - \cos\chi\sin\psi\,\mathbf{e}_3, \tag{S4.2a}$$

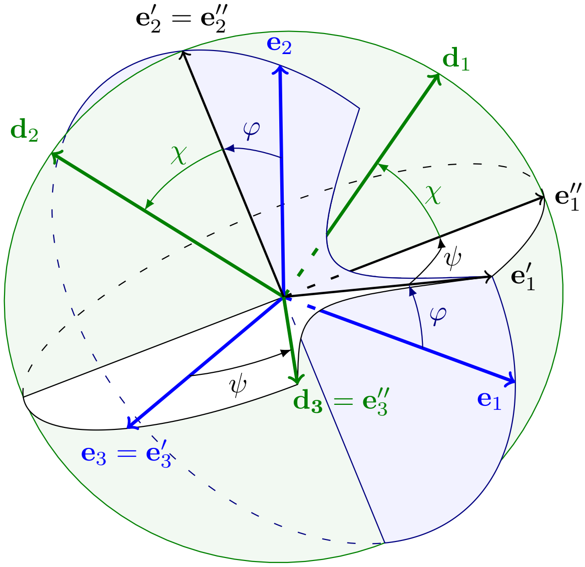

Figure S3: The relationship of the directors $\{\mathbf{d}_j\}$ to the fixed basis $\{\mathbf{e}_j\}$ via the Euler angles $\chi$, $\psi$ and $\varphi$.

$$\mathbf{d}_2 = -\left(\sin\chi\cos\psi\cos\varphi + \cos\chi\sin\varphi\right)\mathbf{e}_1 - \left(\sin\chi\cos\psi\sin\varphi - \cos\chi\cos\varphi\right)\mathbf{e}_2 + \sin\chi\sin\psi\,\mathbf{e}_3, \tag{S4.2b}$$

$$\mathbf{d}_3 = \sin\psi\cos\varphi\,\mathbf{e}_1 + \sin\psi\sin\varphi\,\mathbf{e}_2 + \cos\psi\,\mathbf{e}_3. \tag{S4.2c}$$

Consequently, the strains can be written as

$$u_1 = \frac{\partial\psi}{\partial s}\sin\chi - \frac{\partial\varphi}{\partial s}\cos\chi\sin\psi, \quad u_2 = \frac{\partial\psi}{\partial s}\cos\chi + \frac{\partial\varphi}{\partial s}\sin\chi\sin\psi \quad \text{and} \quad u_3 = \frac{\partial\chi}{\partial s} + \frac{\partial\varphi}{\partial s}\cos\psi. \tag{S4.3}$$

Moreover, denoting by $m_j$ the components of the resultant moment with respect to the fixed basis $\{\mathbf{e}_j\}$, the constitutive assumption (2.8) in the main text leads to

$$\begin{aligned} m_1 =& EI\left\{\cos\psi\cos\varphi\left[(u_1 - u_1^\star)\cos\chi - (u_2 - u_2^\star)\sin\chi\right]\right. \\ &\left. - \sin\varphi\left[(u_1 - u_1^\star)\sin\chi + (u_2 - u_2^\star)\cos\chi\right]\right\} + \mu J\,(u_3 - u_3^\star)\sin\psi\cos\varphi, \end{aligned} \tag{S4.4a}$$

$$\begin{aligned} m_2 =& EI\left\{\cos\psi\sin\varphi\left[(u_1 - u_1^\star)\cos\chi - (u_2 - u_2^\star)\sin\chi\right]\right. \\ &\left. + \cos\varphi\left[(u_1 - u_1^\star)\sin\chi + (u_2 - u_2^\star)\cos\chi\right]\right\} + \mu J\,(u_3 - u_3^\star)\sin\psi\sin\varphi, \end{aligned} \tag{S4.4b}$$

$$m_3 = -EI\sin\psi\left[(u_1 - u_1^\star)\cos\chi - (u_2 - u_2^\star)\sin\chi\right] + \mu J\,(u_3 - u_3^\star)\cos\psi. \tag{S4.4c}$$

##### S4.3 Summary of the reduced model

Under additional hypotheses on the times scales stated in Section 3.1 of the main text, we reduce (S4.1) to get a model suitable for a theoretical study on circumnutations. By focusing on short time periods compared to $\tau_g$ and $\tau_\ell$, we assume a shoot of constant length $\ell$ with constant Young's modulus $E$, whose evolution is governed by

$$\mathbf{m}'(s,t) = -q\,(\ell - s)\,\mathbf{e}_2 \times \mathbf{d}_3(s,t), \tag{S4.5a}$$

$$\dot{u}_1^\star(s,t) = \frac{\alpha}{r\tau_g}\cos\left(\frac{2\pi t}{\tau_e}\right) - \frac{\beta}{r\tau_m\tau_g}\int_{-\infty}^{t-\tau_r} e^{-\frac{1}{\tau_m}(t-\tau_r-\tau)} d_{22}(s,\tau)\,\mathrm{d}\tau - \frac{\eta}{\bar{\tau}_m\tau_g}\int_{-\infty}^{t-\bar{\tau}_r} e^{-\frac{1}{\bar{\tau}_m}(t-\bar{\tau}_r-\tau)} u_1(s,\tau)\,\mathrm{d}\tau, \tag{S4.5b}$$

$$\dot{u}_2^\star(s,t) = \frac{\alpha}{r\tau_g}\sin\left(\frac{2\pi t}{\tau_e}\right) + \frac{\beta}{r\tau_m\tau_g}\int_{-\infty}^{t-\tau_r} e^{-\frac{1}{\tau_m}(t-\tau_r-\tau)} d_{12}(s,\tau)\,\mathrm{d}\tau - \frac{\eta}{\bar{\tau}_m\tau_g}\int_{-\infty}^{t-\bar{\tau}_r} e^{-\frac{1}{\bar{\tau}_m}(t-\bar{\tau}_r-\tau)} u_2(s,\tau)\,\mathrm{d}\tau, \tag{S4.5c}$$

where $d_{ij} := \mathbf{d}_i \cdot \mathbf{e}_j$. Then, in terms of the Euler angles introduced in Section S4.2, we get

$$m_1'(s,t) = -q\,(\ell - s)\cos\psi(s,t), \tag{S4.6a}$$

$$m_2'(s,t) = 0, \tag{S4.6b}$$

$$m_3'(s,t) = q\,(\ell - s)\sin\psi(s,t)\cos\varphi(s,t), \tag{S4.6c}$$

$$\dot u_1^\star(s,t) = \frac{\alpha}{r\tau_g}\cos\left(\frac{2\pi t}{\tau_e}\right) + \frac{\beta}{r\tau_m\tau_g}\int_{-\infty}^{t-\tau_r} e^{-\frac{1}{\tau_m}(t-\tau_r-\tau)}\left(\sin\chi\cos\psi\sin\varphi - \cos\chi\cos\varphi\right)\big|_{(s,\tau)}\,\mathrm{d}\tau$$
$$- \frac{\eta}{\bar\tau_m\tau_g}\int_{-\infty}^{t-\bar\tau_r} e^{-\frac{1}{\bar\tau_m}(t-\bar\tau_r-\tau)}\left[\psi'\sin\chi - \varphi'\cos\chi\sin\psi\right]\big|_{(s,\tau)}\,\mathrm{d}\tau, \tag{S4.6d}$$

$$\dot u_2^\star(s,t) = \frac{\alpha}{r\tau_g}\sin\left(\frac{2\pi t}{\tau_e}\right) + \frac{\beta}{r\tau_m\tau_g}\int_{-\infty}^{t-\tau_r} e^{-\frac{1}{\tau_m}(t-\tau_r-\tau)}\left(\cos\chi\cos\psi\sin\varphi + \sin\chi\cos\varphi\right)\big|_{(s,\tau)}\,\mathrm{d}\tau$$
$$- \frac{\eta}{\bar\tau_m\tau_g}\int_{-\infty}^{t-\bar\tau_r} e^{-\frac{1}{\bar\tau_m}(t-\bar\tau_r-\tau)}\left[\psi'\cos\chi + \varphi'\sin\chi\sin\psi\right]\big|_{(s,\tau)}\,\mathrm{d}\tau, \tag{S4.6e}$$

to be solved for appropriate boundary conditions and initial data.

In the following we explore the contribution of gravitropic and proprioceptive responses.

##### S4.3.1 Graviceptive model: $\alpha = \eta = 0$ and $\beta > 0$

Let us consider the rod model (S4.6) for $\alpha = \eta = 0$. We notice that the planar case studied in [1] is recovered by confining the rod to the $(\mathbf{e}_1, \mathbf{e}_2)$-plane (*i.e.*, $\psi(s,t) = \pi/2$ and $\chi(s,t) = 0$ for all $s$ and $t$), and writing the governing equations in terms of the angle $\theta := \pi/2 - \varphi$. In this case the model suffers an instability and exhibits the onset of a limit cycle as the shoot's length exceeds a critical value. Guided by this result, we extend the analysis to the three-dimensional case.

The steady state solution to problem (S4.6) is given by

$$\chi \equiv 0, \quad \varphi \equiv \frac{\pi}{2}, \quad \psi \equiv \frac{\pi}{2}, \quad u_1^\star \equiv 0, \quad u_2^\star \equiv 0, \tag{S4.7}$$

which corresponds to the straight position along $\mathbf{e}_2$, the axis of gravity. By assuming sufficient regularity, we take the time derivative of equations (S4.6d) and (S4.6e), and we linearize the problem about the equilibrium solution (S4.7), arriving at

$$EI\left(\psi'(s,t) - u_2^\star(s,t)\right)' = -\,q\,(\ell - s)\left(\psi(s,t) - \frac{\pi}{2}\right), \tag{S4.8a}$$
$$\chi''(s,t) = 0, \tag{S4.8b}$$
$$EI\left(\varphi'(s,t) + u_1^\star(s,t)\right)' = -\,q\,(\ell - s)\,,\left(\varphi(s,t) - \frac{\pi}{2}\right), \tag{S4.8c}$$
$$\ddot u_1^\star(s,t) = -\frac{1}{\tau_m}\dot u_1^\star(s,t) + \frac{\beta}{r\tau_g\tau_m}\left(\varphi(s,t-\tau_r) - \frac{\pi}{2}\right), \tag{S4.8d}$$
$$\ddot u_2^\star(s,t) = -\frac{1}{\tau_m}\dot u_2^\star(s,t) - \frac{\beta}{r\tau_g\tau_m}\left(\psi(s,t-\tau_r) - \frac{\pi}{2}\right). \tag{S4.8e}$$

This system of equations is supplemented by the following linearized boundary and initial conditions,

$$\psi(0,t) = \frac{\pi}{2}, \qquad \psi'(\ell,t) - u_2^\star(\ell,t) = 0, \tag{S4.9a}$$
$$\chi(0,t) = 0, \qquad \chi'(\ell,t) = 0, \tag{S4.9b}$$
$$\varphi(0,t) = \frac{\pi}{2}, \qquad \varphi'(\ell,t) + u_1^\star(\ell,t) = 0, \tag{S4.9c}$$

holding $\forall\, t > 0$ as the basal end is clamped and the apical end is torque free, and

$$\varphi(s,t) = \varphi_0(s,t), \qquad \psi(s,t) = \psi_0(s,t), \tag{S4.10a}$$
$$u_1^\star(s,0) = u_{1,0}^\star(s), \qquad u_2^\star(s,0) = u_{2,0}^\star(s), \tag{S4.10b}$$

prescribing respectively the past history of the angular coordinates and the initial datum for the evolution of the spontaneous strains for all $s \in [0,\ell]$.
As for the angle $\chi$, equations (S4.8b) and (S4.9b) yield $\chi(s,t) = 0$ for all $s$ and $t$. Moreover, by assuming sufficient regularity, we can combine the time-derivatives of (S4.8a) and (S4.8c) with the space-derivatives of (S4.8e) and (S4.8d) respectively, so that we get

$$\ddot\psi''(s,t) + \frac{1}{\tau_m}\dot\psi''(s,t) + \frac{q(\ell-s)}{K_1}\left(\ddot\psi(s,t) + \frac{1}{\tau_m}\dot\psi(s,t)\right) + \frac{\beta}{r\tau_m\tau_g}\psi'(s,t-\tau_r) = 0, \tag{S4.11a}$$

$$\ddot{\varphi}''(s,t) + \frac{1}{\tau_m}\dot{\varphi}''(s,t) + \frac{q(\ell - s)}{K_1}\left(\ddot{\varphi}(s,t) + \frac{1}{\tau_m}\dot{\varphi}(s,t)\right) + \frac{\beta}{r\tau_m\tau_g}\varphi'(s,t-\tau_r) = 0, \tag{S4.11b}$$

along with the boundary conditions (S4.9a)$_1$, (S4.9c)$_1$ and

$$\ddot{\psi}'(\ell,t) + \frac{1}{\tau_m}\dot{\psi}'(\ell,t) + \frac{\beta}{r\tau_m\tau_g}\left(\psi(\ell,t-\tau_r) - \frac{\pi}{2}\right) = 0, \tag{S4.12a}$$

$$\ddot{\varphi}'(\ell,t) + \frac{1}{\tau_m}\dot{\varphi}'(\ell,t) + \frac{\beta}{r\tau_m\tau_g}\left(\varphi(\ell,t-\tau_r) - \frac{\pi}{2}\right) = 0, \tag{S4.12b}$$

holding $\forall\, t > 0$ and resulting from time differentiation of (S4.9a)$_2$ and (S4.9c)$_2$.

We notice that equations (S4.11) are decoupled and, up to a shift by $\pi/2$, they are equivalent to the linearization of the planar model about the equilibrium of $\theta \equiv 0$ [1]. Therefore we can rely on the analysis carried out for the planar model to conclude that the trivial equilibrium becomes unstable when the same critical length is attained.

##### S4.3.2 Microgravity: $\alpha = \beta = 0$, $\eta > 0$, and $q = 0$

For $\beta = 0$ and $q = 0$, virtual and current configuration coincide (*i.e.*, $u_j^\star = u_j$ for all $j$), so that equations (S4.5) reduce to

$$\dot{u}_j(s,t) = -\frac{\eta}{\bar{\tau}_m\tau_g}\int_{-\infty}^{t-\bar{\tau}_r} e^{-\frac{1}{\bar{\tau}_m}(t-\bar{\tau}_r-\tau)}u_j(s,\tau)\,\mathrm{d}\tau, \tag{S4.13}$$

for $j = 1, 2$. By assuming sufficient regularity, a time differentiation yields

$$\ddot{u}_j(s,t) + \frac{1}{\bar{\tau}_m}\dot{u}_j(s,t) + \frac{\eta}{r\tau_g\bar{\tau}_m}u_j(s,t-\bar{\tau}_r) = 0, \tag{S4.14}$$

for $j = 1, 2$, which can be restated in dimensionless form as

$$\ddot{\hat{u}}_j(\hat{s},\hat{t}) + \frac{\bar{\tau}_r}{\bar{\tau}_m}\dot{\hat{u}}_j(\hat{s},\hat{t}) + \frac{\eta\,\bar{\tau}_r^2}{\bar{\tau}_m\tau_g}\hat{u}_j(\hat{s},\hat{t}-1) = 0, \tag{S4.15}$$

where $\hat{u}_j(\hat{s},\hat{t}) := u_j(\hat{s}\ell,\hat{t}\tau_r)$ for $j = 1, 2$, and dots and primes denote differentiation with respect to $\hat{t} := t/\bar{\tau}_r$ and $\hat{s} := s/\ell$, respectively.

Since equation (S4.15) does not contain space derivatives, we can rely on the theory of retarded functional differential equations (RFDEs) by considering the space variable as a parameter. Indeed, given the problem

$$\ddot{u}(s,t) + a\dot{u}(s,t) + bu(s,t-1) = 0, \qquad s \in [0,\ell_0]\,,\ t > 1, \tag{S4.16a}$$

$$u(s,t) = u_0(s,t), \qquad s \in [0,\ell_0]\,,\ t \in [0,1]\,, \tag{S4.16b}$$

with $a > 0$ and an initial datum $u_0$ that is regular enough, say $u_0 \in C^\infty$, we can consider the solution $u(s,t) := u_s(t)$ where $u_s(t)$ is the unique solution to

$$\ddot{v}(t) + a\dot{v}(t) + bv(t-1) = 0, \qquad t > 1, \tag{S4.17a}$$

$$v(t) = u_0(s,t), \qquad t \in [0,1]\,, \tag{S4.17b}$$

for any fixed $s \in [0,\ell_0]$. Then the regularity of $u(s,t)$ with respect to $s$ follows from the results on the continuous dependence of solutions to RFDEs on initial data [8]. Moreover, we can exploit the stability analysis of the trivial equilibrium of (S4.17) to learn something about the solution $u(s,t)$. To this aim, we restate the following lemma that holds for the characteristic equation associated with (S4.17), see [1, 15].

**Lemma S4.1.** *Consider the equation*

$$\left(\omega^2 + a\omega\right)e^\omega + b = 0, \tag{S4.18}$$

*for $b > 0$, and let $\xi_b$ be the unique solution of $\xi^2 = b\cos(\xi)$ in $(0,\pi/2)$ and $a_b := \sin(\xi_b)b/\xi_b$. Then the following holds for equation* (S4.18)*:*

*(i) All roots have negative real parts if and only if $a > a_b$;*

*(ii) For $a = a_b$, $\pm i\xi_b$ is the only pair of simple imaginary roots. In particular, no other root is an integer multiple of $i\xi_b$;*

*(iii) There exists an $\epsilon > 0$ and a root $\omega(a)$ that is continuously differentiable in $(a_b - \epsilon, a_b + \epsilon)$ s.t. $\omega(a_b) = i\xi_b$ and $\mathrm{Re}\,(\omega'(a_b)) < 0$;*

*(iv) For each $a < a_b$, there exist precisely two roots $\omega$ with $\mathrm{Re}\,(\omega) > 0$ and $\mathrm{Im}\,(\omega) \in (-\pi, \pi)$.*

In addition to this, we can show the following.

**Lemma S4.2.** *Consider equation (S4.18) for $b > 0$ and let $b_a := (a + 2\tilde{\omega})\, e^{\tilde{\omega}}$ where $\tilde{\omega} := (\sqrt{4 + a^2} - a - 2)/2$. Then for $b < b_a$ there exist precisely two real roots, which coincide for $b = b_a$, whereas there exist no real roots for $b > b_a$.*

*Proof.* Let us define $y(\omega) := \omega^2 + a\omega$ and $z(\omega) := -be^{-\omega}$. By means of the graphical method, one can show that there are at most two real intersections between the graphs of $y$ and $z$. If there is a single distinct real root $\tilde{\omega}$, then it is such that

$$y(\tilde{\omega}) = z(\tilde{\omega}) \quad \text{and} \quad \frac{\partial y}{\partial \omega}(\tilde{\omega}) = \frac{\partial z}{\partial \omega}(\tilde{\omega}), \tag{S4.19}$$

namely, $\tilde{\omega}^2 + a\tilde{\omega} + be^{-\tilde{\omega}} = 0$ and $2\tilde{\omega} + a = be^{-\tilde{\omega}}$. Therefore $\tilde{\omega}$ needs to solve $\tilde{\omega}^2 + (2+a)\tilde{\omega} + a = 0$, whose solutions are

$$\omega_{\pm} = \frac{-2 - a \pm \sqrt{4 + a^2}}{2}. \tag{S4.20}$$

Since $2\tilde{\omega} + a = be^{-\tilde{\omega}} > 0$, we conclude that the only admissible root is given by $\tilde{\omega} = \omega_+$ and that $b = (a + 2\tilde{\omega})e^{\tilde{\omega}} =: b_a$. □

Finally we can prove the following facts.

**Theorem S4.3.** *Consider problem (S4.16) and the solution $u(s,t) := u_s(t)$ where $u_s$ solves (S4.17) for any $s \in [0, \ell_0]$. Moreover, let $a_b$ be defined as in Lemma S4.1. Then,*

*(i) if $a > a_b$, the trivial equilibrium of (S4.16a) is stable and there exists $\delta > 0$ such that $||u_0||_\infty < \delta$ implies that, for any fixed $s \in [0, \ell_0]$, $|u(s,t)| \to 0$ as $t \to \infty$;*

*(ii) if $a < a_b$, the trivial equilibrium of (S4.16a) is unstable.*

*Moreover, for $a = a_b$, equation (S4.16a) admits nontrivial periodic solutions.*

*Proof.* Assume that $a > a_b$. Then, by means of Lemma S4.1, for any $\epsilon > 0$, there exists $\delta_\epsilon > 0$ such that $\sup |u_0(s,t)| < \delta_\epsilon$ implies $|u_s(t)| < \epsilon$ for all $t \geq 0$. Moreover, there exists $\delta > 0$ such that $\sup |u_0(s,t)| < \delta$ implies that $|u_s(t)| \to 0$ as $t \to \infty$. It follows that if we take $u_0(s,t)$ such that $\sup |u_0(s,t)| < \min\{\delta_\epsilon, \delta\}$, then $|u_s(t)| < \epsilon$ for all $(s,t) \in [0, \ell_0] \times [0, \infty)$ and, for any fixed $s \in [0, \ell_0]$, $|u(s,t)| \to 0$ as $t \to \infty$.
On the contrary, if $a < a_b$, there exists $\epsilon > 0$ such that for any $\delta > 0$, we find an initial datum $\bar{u}(t)$ for which $\sup |\bar{u}(t)| < \delta$ and the corresponding solution of (S4.17a) verifies $|u(t)| > \epsilon$ for some $t \geq 0$. Then the statement follows by observing that $u(s,t) := u(t)$ solves (S4.16) for the space-independent initial datum $u_0(s,t) := \bar{u}(t)$. □

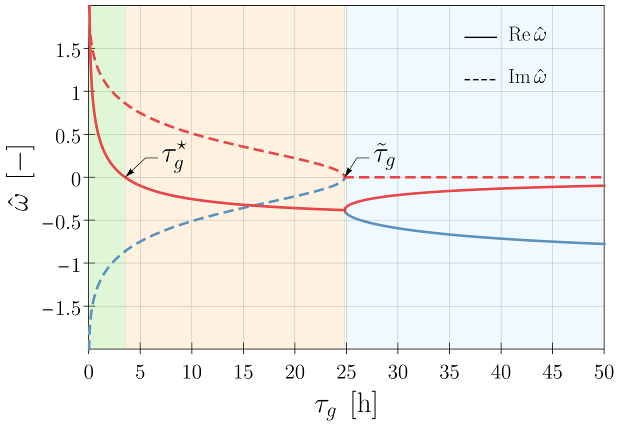

Figure S4: Real (solid lines) and imaginary (dashed lines) part of two roots of the characteristic equation $\hat{\omega}^2 + (\bar{\tau}_r/\bar{\tau}_m)\,\hat{\omega} + [\eta\,\bar{\tau}_r^2/(\bar{\tau}_m\tau_g)]\, e^{-\hat{\omega}} = 0$, as functions of $\tau_g \in [0, 50]$ h and for $\eta = 20$, $\bar{\tau}_r = \bar{\tau}_m = 12$ min, $r = 0.5 \times 10^{-3}$ m. We distinguish three regions corresponding to different dynamical responses: (i) an exponential decay for $\tau_g > \tilde{\tau}_g \approx 24.83$ h (light blue), (ii) a damped oscillation (orange), and (iii) an increasing oscillation for $\tau_g < \tau_g^\star \approx 3.52$ h (green).

Finally, by applying the results above to equation (S4.15), we conclude that it admits nontrivial periodic solutions when

$$\tau_g = \tau_g^\star := \eta\bar{\tau}_r \frac{\sin(\xi^\star)}{\xi^\star}, \tag{S4.21}$$

where $\xi^\star$ is the unique solution of $\xi \tan\xi = \bar{\tau}_r/\bar{\tau}_m$ in $(0, \pi/2)$. Moreover, the characteristic equation associated with (S4.15) has a pair of conjugate complex roots for

$$\tau_g < \tilde{\tau}_g := \frac{\eta\bar{\tau}_r^2 e^{-\omega_0}}{2\omega_0\bar{\tau}_m + \bar{\tau}_r}, \tag{S4.22}$$

where $\omega_0 := \left(1 + \bar{\tau}_r^2/(4\bar{\tau}_m^2)\right)^{1/2} - (1 + \bar{\tau}_r/(2\bar{\tau}_m))$, and their real part crosses zero at $\tau_g^\star$, as shown in Fig. S4. Then the trivial straight position is stable for $\tau_g > \tau_g^\star$ and unstable for $\tau_g < \tau_g^\star$. Therefore, when the growth velocity is sufficiently fast, our model exhibits oscillations about the equilibrium configuration also under microgravity conditions, regardless of the shoot length. However, we notice that for the present choice of model parameters, the critical value $\tau_g^\star$ is one order of magnitude smaller than the observed growth times, cf. Table 1 in the main text.

##### S4.3.3 Proprio-graviceptive model: $\alpha = 0$ and $\beta, \eta > 0$

As regards the proprio-graviceptive model corresponding to $\alpha = 0$ and $\beta$, $\eta > 0$, we first obtain planar steady-state solutions, which might be useful to estimate the ratio between gravitropic and proprioceptive sensitivities from experimental observations, and then we carry out the stability analysis as done for the graviceptive case.

**Planar steady-state solutions.** By confining the rod model (S4.6) to the plane $(\mathbf{e}_1, \mathbf{e}_2)$, and assuming sufficient regularity, we get

$$\begin{aligned} K_1\left[\theta'(s,t) - u_1^\star(s,t)\right]' &= -\, q(\ell - s)\sin\theta(s,t), && \text{(S4.23a)}\\ r\,\tau_g \dot{u}_1^\star(s,t) &= -\,\beta w_g(s,t) - r\,\eta\, w_p(s,t), && \text{(S4.23b)}\\ \tau_m \dot{w}_g(s,t) &= -\, w_g(s,t) + \sin\theta(s, t-\tau_r), && \text{(S4.23c)}\\ \bar{\tau}_m \dot{w}_p(s,t) &= -\, w_p(s,t) + \theta'(s, t-\tau_r), && \text{(S4.23d)} \end{aligned}$$

as the governing equations, where

$$w_g := \frac{1}{\tau_m}\int_{-\infty}^{t-\tau_r} e^{-\frac{1}{\tau_m}(t-\tau_r-\tau)} \sin\theta(s,\tau)\,\mathrm{d}\tau \quad \text{and} \quad w_p := \frac{1}{\bar{\tau}_m}\int_{-\infty}^{t-\bar{\tau}_r} e^{-\frac{1}{\bar{\tau}_m}(t-\bar{\tau}_r-\tau)}\theta'(s,\tau)\,\mathrm{d}\tau. \tag{S4.24}$$

Then a steady-state solution $\theta(s)$ of (S4.23) needs to solve

$$\hat{\theta}'(\hat{s}) = -\frac{\beta}{\eta}\frac{\ell}{r}\sin\hat{\theta}(\hat{s}), \tag{S4.25}$$

for $\hat{s} \in [0, 1]$, combined with the boundary condition $\hat{\theta}(0) = \theta_0$. Here, $\hat{\theta}(\hat{s}) := \theta(\hat{s}\ell)$ and primes denote differentiation with respect to $\hat{s} := s/\ell$. Then an equilibrium of (S4.23) is given by

$$\hat{\theta}(\hat{s}) = 2\,\mathrm{acot}\left[\cot\left(\tfrac{\theta_0}{2}\right) e^{\frac{\beta}{\eta}\frac{\ell}{r}\hat{s}}\right], \tag{S4.26}$$

for $\hat{s} \in [0, 1]$. Therefore, when converging to (S4.26), the final shape is completely determined by the ratio between the two sensitivities, $\beta/\eta$, while the whole dynamics towards the steady state depends also on the characteristic times, *i.e.*, $\tau_g$, $\tau_m$, $\tau_r$, $\bar{\tau}_m$ and $\bar{\tau}_r$. Since gravitropic and proprioceptive responses generate planar dynamics for initially straight plant shoots, the planar steady-state solution (S4.26) can be used to determine the dimensionless parameter $\beta\ell/(\eta r)$ by fitting the experimental shapes attained in a time period that is short with respect to growth, as already done for the instantaneous version of this model without gravity loads [2].

**Stability analysis.** Guided by the analysis carried out in Section S4.3.1, we linearize (S4.6) about the equilibrium (S4.7), thus arriving at

$$\begin{aligned} \psi''(s,t) &= u_2^{\star\prime}(s,t) - \frac{q}{EI}(\ell - s)\left(\psi(s,t) - \frac{\pi}{2}\right), && \text{(S4.27a)}\\ \chi''(s,t) &= 0, && \text{(S4.27b)}\\ \varphi''(s,t) &= -\, u_1^{\star\prime}(s,t) - \frac{q}{EI}(\ell - s)\left(\varphi(s,t) - \frac{\pi}{2}\right), && \text{(S4.27c)} \end{aligned}$$

$$\dot{u}_1^{\star}(s,t) = \frac{\beta}{r\tau_m\tau_g}\int_{-\infty}^{t-\tau_r} e^{-\frac{1}{\tau_m}(t-\tau_r-\tau)}\left(\varphi(s,\tau)-\frac{\pi}{2}\right)\mathrm{d}\tau + \frac{\eta}{\bar{\tau}_m\tau_g}\int_{-\infty}^{t-\bar{\tau}_r} e^{-\frac{1}{\bar{\tau}_m}(t-\bar{\tau}_r-\tau)}\varphi'(s,\tau)\,\mathrm{d}\tau, \tag{S4.27d}$$

$$\dot{u}_2^{\star}(s,t) = -\frac{\beta}{r\tau_m\tau_g}\int_{-\infty}^{t-\tau_r} e^{-\frac{1}{\tau_m}(t-\tau_r-\tau)}\left(\psi(s,\tau)-\frac{\pi}{2}\right)\mathrm{d}\tau - \frac{\eta}{\bar{\tau}_m\tau_g}\int_{-\infty}^{t-\bar{\tau}_r} e^{-\frac{1}{\bar{\tau}_m}(t-\bar{\tau}_r-\tau)}\psi'(s,\tau)\,\mathrm{d}\tau. \tag{S4.27e}$$

By assuming sufficient regularity, we get

$$\dot{\psi}''(s,t) + \frac{\beta}{r\tau_m\tau_g}\int_{-\infty}^{t-\tau_r} e^{-\frac{1}{\tau_m}(t-\tau_r-\tau)}\psi'(s,\tau)\,\mathrm{d}\tau + \frac{\eta}{\bar{\tau}_m\tau_g}\int_{-\infty}^{t-\bar{\tau}_r} e^{-\frac{1}{\bar{\tau}_m}(t-\bar{\tau}_r-\tau)}\psi''(s,\tau)\,\mathrm{d}\tau + \frac{q}{EI}(\ell-s)\,\dot{\psi}(s,t) = 0, \tag{S4.28a}$$

$$\dot{\varphi}''(s,t) + \frac{\beta}{r\tau_m\tau_g}\int_{-\infty}^{t-\tau_r} e^{-\frac{1}{\tau_m}(t-\tau_r-\tau)}\varphi'(s,\tau)\,\mathrm{d}\tau + \frac{\eta}{\bar{\tau}_m\tau_g}\int_{-\infty}^{t-\bar{\tau}_r} e^{-\frac{1}{\bar{\tau}_m}(t-\bar{\tau}_r-\tau)}\varphi''(s,\tau)\,\mathrm{d}\tau + \frac{q}{EI}(\ell-s)\,\dot{\varphi}(s,t) = 0, \tag{S4.28b}$$

where equations (S4.27d) and (S4.27e) have been combined with the time derivatives of equations (S4.27a) and (S4.27c), respectively, while solving equation (S4.27b) with boundary conditions (S4.9b). We notice that equations (S4.28) along with the boundary conditions $(\text{S4.9a})_1$, $(\text{S4.9c})_1$ and (S4.12), form two equivalent decoupled problems. Moreover, up to a shift of $\pi/2$, such a problem is precisely the linearization of the planar model (S4.23) about the trivial solution $\theta \equiv 0$, which can be restated in dimensionless form as

$$\dot{\hat{\theta}}''(\hat{s},\hat{t}) + \beta\frac{\ell}{r}\frac{\tau_s^2}{\tau_m\tau_g}\int_{-\infty}^{\hat{t}-\frac{\tau_r}{\tau_s}} e^{-\frac{\tau_s}{\tau_m}\left(\hat{t}-\frac{\tau_r}{\tau_s}-\tau\right)}\hat{\theta}'(\hat{s},\tau)\,\mathrm{d}\tau + \eta\frac{\tau_s^2}{\bar{\tau}_m\tau_g}\int_{-\infty}^{\hat{t}-\frac{\bar{\tau}_r}{\tau_s}} e^{-\frac{\tau_s}{\bar{\tau}_m}\left(\hat{t}-\frac{\bar{\tau}_r}{\tau_s}-\tau\right)}\hat{\theta}''(\hat{s},\tau)\,\mathrm{d}\tau + \frac{q\ell^3}{K_1}(1-\hat{s})\dot{\hat{\theta}}(\hat{s},\hat{t}) = 0, \tag{S4.29}$$

with boundary conditions

$$\hat{\theta}(0,\hat{t}) = 0, \tag{S4.30a}$$

$$\frac{\tau_g}{\tau_s^2}\dot{\hat{\theta}}'(1,\hat{t}) = -\frac{\beta}{\tau_m}\frac{\ell}{r}\int_{-\infty}^{\hat{t}-\frac{\tau_r}{\tau_s}} e^{-\frac{\tau_s}{\tau_m}\left(\hat{t}-\frac{\tau_r}{\tau_s}-\tau\right)}\hat{\theta}(1,\tau)\,\mathrm{d}\tau - \frac{\eta}{\bar{\tau}_m}\int_{-\infty}^{\hat{t}-\frac{\bar{\tau}_r}{\tau_s}} e^{-\frac{\tau_s}{\bar{\tau}_m}\left(\hat{t}-\frac{\bar{\tau}_r}{\tau_s}-\tau\right)}\hat{\theta}'(1,\tau)\,\mathrm{d}\tau, \tag{S4.30b}$$

for $\hat{t} > 0$, where $\hat{\theta}(\hat{s},\hat{t}) := \theta(\hat{s}\ell,\hat{t}\tau_s)$ for any fixed time scale $\tau_s$. By seeking time-harmonic solutions of the form $\hat{\theta}(\hat{s},\hat{t}) = \Theta(\hat{s})e^{\hat{\omega}\hat{t}}$ with $\mathrm{Re}(\hat{\omega}) > -\min\{\tau_s/\tau_m, \tau_s/\bar{\tau}_m\}$, we get

$$a\Theta''(\hat{s}) + b\Theta'(\hat{s}) + c(1-\hat{s})\Theta(\hat{s}) = 0, \tag{S4.31}$$

where

$$a := \hat{\omega} + \eta\frac{\tau_s^2 e^{-\hat{\omega}\frac{\bar{\tau}_r}{\tau_s}}}{\tau_g(\bar{\tau}_m\hat{\omega}+\tau_s)}, \qquad b := \beta\frac{\ell}{r}\frac{\tau_s^2 e^{-\hat{\omega}\frac{\tau_r}{\tau_s}}}{\tau_g(\tau_m\hat{\omega}+\tau_s)}, \qquad \text{and} \qquad c := \hat{\omega}\frac{q\ell^3}{K_1}. \tag{S4.32}$$

By imposing the boundary conditions (S4.30), and neglecting the trivial case, we derive the following characteristic equation for the dimensionless frequency $\hat{\omega}$, namely,

$$\mathrm{Ai}(x_0)\left[b\,\mathrm{Bi}(x_1) + 2\sqrt[3]{a^2c}\,\mathrm{Bi}'(x_1)\right] - \mathrm{Bi}(x_0)\left[b\,\mathrm{Ai}(x_1) + 2\sqrt[3]{a^2c}\,\mathrm{Ai}'(x_1)\right] = 0, \tag{S4.33}$$

where

$$x_0 := \frac{b^2-4ac}{4a\sqrt[3]{ac^2}}, \quad x_1 := \frac{b^2}{4a\sqrt[3]{ac^2}}, \tag{S4.34}$$

and $\mathrm{Ai}(x)$, $\mathrm{Bi}(x)$ are the Airy functions of the first and second kind, respectively, and a prime denotes differentiation of such functions with respect to their argument.

Then we can explore the stability of the model by performing a numerical study of the roots of equation (S4.33) for shoots of increasing length $\ell$. To this aim, we exploited the FindRoot functionality of Mathematica v12.0.0.0. In agreement with the literature, we calibrated the model by setting $\beta = 0.8$, $\tau_g = 20\,\mathrm{h}$, $\tau_m = \tau_r = 12\,\mathrm{min}$, $r = 0.5\,\mathrm{mm}$, $E = 10^7\,\mathrm{Pa}$ for the Young's modulus, and $\rho = 10^3\,\mathrm{Kg\,m^{-3}}$ for the mass density. As for $\eta$, $\bar{\tau}_r$ and $\bar{\tau}_m$, we estimated their order of magnitude by qualitatively fitting the steady-state solution (S4.26) and the dynamics reported in [12], and

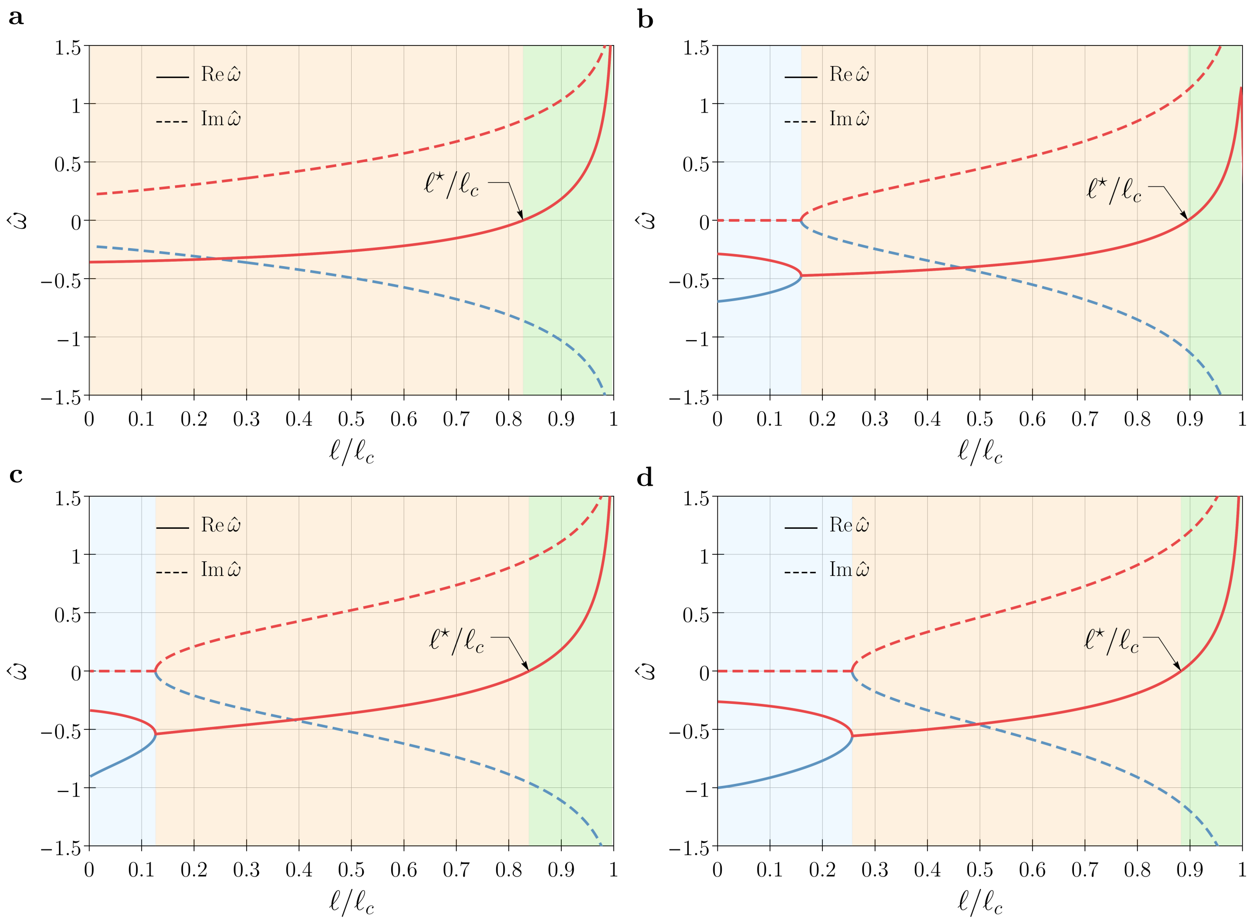

Figure S5: Real (solid lines) and imaginary (dashed lines) part of the roots of the characteristic equation (S4.33) as functions of $\ell/\ell_c \in [0, 1]$ and for the model parameters introduced in the section. We distinguish three regions corresponding to different dynamical responses: (i) an exponential decay (light blue), (ii) a damped oscillation (orange), and (iii) an increasing oscillation (green) for $\ell > \ell^\star$. More specifically, we get (a) $\ell^\star \approx 0.827\,\ell_c$ for $\bar\tau_r = \bar\tau_m = 12$ min, (b) $\ell^\star \approx 0.896\,\ell_c$ for $\bar\tau_r = 1$ min and $\bar\tau_m = 12$ min, (c) $\ell^\star \approx 0.838\,\ell_c$ for $\bar\tau_r = 12$ min and $\bar\tau_m = 6$ min, and (d) $\ell^\star \approx 0.883\,\ell_c$ for $\bar\tau_r = 6$ min and $\bar\tau_m = 6$ min. For comparison, the same choice of model parameters yield a critical value of $\ell^\star \approx 0.895\,\ell_c$ in the gravitropic case [1].

we explored values around $\eta = 20$ and $\bar\tau_m = \bar\tau_r = 12\,\text{min}$. For such a choice of model parameters, we determined the values of the frequency $\hat\omega$ letting $\ell$ range in $[0, \ell_c]$. Here, $\ell_c$ denotes the critical length at which an elastic rod of bending stiffness $EI$ subject to a distributed vertical load of magnitude $q$ loses stability, *i.e.*,

$$\ell_c := \sqrt[3]{\alpha_0 EI/q}, \tag{S4.35}$$

with $\alpha_0 \approx 7.837$, see [7].

Fig. S5 shows the real (solid lines) and the imaginary (dashed lines) part of two roots of (S4.33). As for the case of the gravitropic rod model [1], we distinguish in the figure three regions corresponding to different dynamical responses: (i) an exponential decay (light blue region, where roots are real and negative), (ii) a damped oscillation (orange region, where roots are complex conjugate with negative real part), and (iii) an increasing oscillation (green region, where roots are complex conjugate with positive real part) for $\ell > \ell^\star$. We remark the fact that the memory time $\bar\tau_m$ and the delay $\bar\tau_r$ influence the value of the critical length $\ell^\star$: Lower times $\bar\tau_r$ and $\bar\tau_m$ imply a lower critical length. This affects the overall effect of the proprioceptive term, which can either destabilize (Fig. S5a,c,d) or stabilize (Fig. S5b) the system with respect to the gravitropic case ($\eta = 0$).

#### S5 Computational model

The nonlinear response of (S4.1) and (S4.5) have been explored by a computational model implemented in a Python3 code that exploits the DOLFIN library as interface for the FEniCS Project Version 2019.1.0 [10]. Since the numerical scheme for the full model (S4.1) can be easily adapted to the reduced version given by (S4.5), in the following we deal only with the former.

An effective way to implement the model is to write all equations in the reference domain $\mathcal{B}_0$, *i.e.*, in terms of the parameter $S \in [0, \ell_0]$. Any material field can be converted into a spatial field, and vice versa. Indeed, as shown in Supplemental Section S1, the motion $s(S,t)$ can be analytically determined for the growth law given by (S4.1b), namely,

$$s(S,t) = \begin{cases} S & \text{if } S \leq \ell_0 - \ell_g, \\ \ell(t^\star(S)) - \ell_g & \text{if } S > \ell_0 - \ell_g \text{ and } t \geq t^\star(S), \\ \ell(t) - (\ell_0 - S)e^{t/\tau_g} & \text{if } S > \ell_0 - \ell_g \text{ and } t < t^\star(S), \end{cases} \tag{S5.1}$$

where

$$\ell(t) = \begin{cases} \ell_0 e^{t/\tau_g} & \text{if } t \leq t^\star(0), \\ \max\{\ell_0, \ell_g\} + \frac{\ell_g}{\tau_g}\left(t - \max\{0, t^\star(0)\}\right) & \text{if } t > t^\star(0), \end{cases} \quad \text{and} \quad t^\star(S) = \tau_g \ln\left(\frac{\ell_g}{\ell_0 - S}\right). \tag{S5.2}$$

Moreover, its inverse is given by

$$S(s,t) := \begin{cases} s & \text{if } s \leq \ell_0 - \ell_g, \\ \ell_0 + \left[s - \ell\left(\bar{t}^\star(s,t)\right)\right] e^{-\bar{t}^\star(s,t)/\tau_g} & \text{if } s \in (\ell_0 - \ell_g, \ell(t) - \ell_g], \\ \ell_0 + \left[s - \ell(t)\right] e^{-t/\tau_g} & \text{if } s \in (\ell(t) - \ell_g, \ell(t)], \end{cases} \tag{S5.3}$$

where $\bar{t}^\star(s,t) := t + \tau_g\left(s + \ell_g - \ell(t)\right)/\ell_g$.
As a first step towards the numerical formulation, we introduce some auxiliary fields representing the delay integrals, namely,

$$w_{1,g}(S,t) := -\frac{1}{\tau_m}\int_{-\infty}^{t-\tau_r} e^{-\frac{1}{\tau_m}(t-\tau_r-\tau)} \sin\theta_h(S,\tau)\cos\alpha_h(S,\tau)\,\mathrm{d}\tau, \tag{S5.4a}$$

$$w_{1,p}(S,t) := -\frac{1}{\bar{\tau}_m}\int_{-\infty}^{t-\bar{\tau}_r} e^{-\frac{1}{\bar{\tau}_m}(t-\bar{\tau}_r-\tau)} u_1(S,\tau)\,\mathrm{d}\tau, \tag{S5.4b}$$

$$w_{2,g}(S,t) := \frac{1}{\tau_m}\int_{-\infty}^{t-\tau_r} e^{-\frac{1}{\tau_m}(t-\tau_r-\tau)} \cos\theta_h(S,\tau)\,\mathrm{d}\tau, \tag{S5.4c}$$

$$w_{2,p}(S,t) := -\frac{1}{\bar{\tau}_m}\int_{-\infty}^{t-\bar{\tau}_r} e^{-\frac{1}{\bar{\tau}_m}(t-\bar{\tau}_r-\tau)} u_2(S,\tau)\,\mathrm{d}\tau, \tag{S5.4d}$$

so that such integrals may be computed from the solution of the following differential equations

$$\frac{\mathrm{d}w_{1,g}}{\mathrm{d}t} = -\frac{1}{\tau_m}w_{1,g} - \frac{1}{\tau_m}\sin\theta_h(S,t-\tau_r)\cos\alpha_h(S,t-\tau_r), \tag{S5.5a}$$

$$\frac{\mathrm{d}w_{1,p}}{\mathrm{d}t} = -\frac{1}{\bar{\tau}_m}w_{1,p} - \frac{1}{\bar{\tau}_m}u_1(S,t-\bar{\tau}_r), \tag{S5.5b}$$

$$\frac{\mathrm{d}w_{2,g}}{\mathrm{d}t} = -\frac{1}{\tau_m}w_{2,g} + \frac{1}{\tau_m}\cos\theta_h(S,t-\tau_r), \tag{S5.5c}$$

$$\frac{\mathrm{d}w_{2,p}}{\mathrm{d}t} = -\frac{1}{\bar{\tau}_m}w_{2,p} - \frac{1}{\bar{\tau}_m}u_2(S,t-\bar{\tau}_r), \tag{S5.5d}$$

respectively. Then we can write the governing equations in terms of the Euler angles introduced in S4.2 and the angles describing the statoliths pile configuration as defined in S3, *i.e.*,

$$\frac{\partial m_1}{\partial S} = -\,q\left[\ell(t) - s(S,t)\right]\lambda\cos\psi, \tag{S5.6a}$$

$$\frac{\partial m_2}{\partial S} = 0, \tag{S5.6b}$$

$$\frac{\partial m_3}{\partial S} = q\left[\ell(t) - s(S,t)\right]\lambda\sin\psi\cos\varphi, \tag{S5.6c}$$

$$\begin{aligned}\tau_a \frac{\mathrm{d}\theta_h}{\mathrm{d}t} &= \cos\theta_h \left[\cos\chi\cos\alpha_h\cos\varphi + (-\cos\psi\cos\alpha_h\sin\chi + \sin\psi\sin\alpha_h)\sin\varphi\right] \\ &\quad - (\cos\varphi\sin\chi + \cos\chi\cos\psi\sin\varphi)\sin\theta_h, \qquad \text{(S5.6d)}\\ \tau_a \frac{\mathrm{d}\alpha_h}{\mathrm{d}t}\sin\theta_h &= -\cos\chi\cos\varphi\sin\alpha_h + (\cos\alpha_h\sin\psi + \cos\psi\sin\chi\sin\alpha_h)\sin\varphi, \qquad \text{(S5.6e)}\\ \frac{\mathrm{d}u_1^\star}{\mathrm{d}t} &= \frac{1}{\lambda}\frac{\mathrm{d}\lambda}{\mathrm{d}t}\left[\frac{\alpha}{r}\cos\left(\frac{2\pi t}{\tau_e}\right) + \frac{\beta}{r}w_{1,g} + \eta w_{1,p}\right], \qquad \text{(S5.6f)}\\ \frac{\mathrm{d}u_2^\star}{\mathrm{d}t} &= \frac{1}{\lambda}\frac{\mathrm{d}\lambda}{\mathrm{d}t}\left[\frac{\alpha}{r}\sin\left(\frac{2\pi t}{\tau_e}\right) + \frac{\beta}{r}w_{2,g} + \eta w_{2,p}\right], \qquad \text{(S5.6g)}\end{aligned}$$

where $\lambda(S,t) = \frac{\partial s(S,t)}{\partial S}$ and

$$\begin{aligned} m_1 =& EI\left\{\cos\psi\cos\varphi\left[(u_1-u_1^\star)\cos\chi - (u_2-u_2^\star)\sin\chi\right]\right. \\ & \left. -\sin\varphi\left[(u_1-u_1^\star)\sin\chi + (u_2-u_2^\star)\cos\chi\right]\right\} + \mu J\,(u_3-u_3^\star)\sin\psi\cos\varphi, \qquad \text{(S5.7a)}\\ m_2 =& EI\left\{\cos\psi\sin\varphi\left[(u_1-u_1^\star)\cos\chi - (u_2-u_2^\star)\sin\chi\right]\right. \\ & \left. +\cos\varphi\left[(u_1-u_1^\star)\sin\chi + (u_2-u_2^\star)\cos\chi\right]\right\} + \mu J\,(u_3-u_3^\star)\sin\psi\sin\varphi, \qquad \text{(S5.7b)}\\ m_3 =& -EI\sin\psi\left[(u_1-u_1^\star)\cos\chi - (u_2-u_2^\star)\sin\chi\right] + \mu J\,(u_3-u_3^\star)\cos\psi, \qquad \text{(S5.7c)}\end{aligned}$$

with

$$u_1 = \frac{1}{\lambda}\left[\frac{\partial\psi}{\partial S}\sin\chi - \frac{\partial\varphi}{\partial S}\cos\chi\sin\psi\right],\ u_2 = \frac{1}{\lambda}\left[\frac{\partial\psi}{\partial S}\cos\chi + \frac{\partial\varphi}{\partial S}\sin\chi\sin\psi\right],\ u_3 = \frac{1}{\lambda}\left[\frac{\partial\chi}{\partial S} + \frac{\partial\varphi}{\partial S}\cos\psi\right]. \qquad \text{(S5.8)}$$

The weak formulation of (S5.5)-(S5.6) is obtained by multiplying such equations by the test functions and integrating by parts in space along the interval $[0, \ell_0]$ while accounting for the appropriate boundary conditions. Following standard finite element procedures, the unknowns are discretized in space using linear Langrange shape functions, while for the time discretization we used the backward Euler method. Finally, the rod axis $\mathbf{p}$ can be reconstructed by integrating in space the tangent that, for unshearable rods, coincides with the director $\mathbf{d}_3$, *i.e.*,

$$\mathbf{p}(S,t) = \mathbf{p}(0,t) + \int_0^{\ell_0} \lambda(\zeta,t)\mathbf{d}_3(\zeta,t)\,\mathrm{d}\zeta. \qquad \text{(S5.9)}$$

The Supplemental Videos 1 and 2 provide illustrative examples of the results obtained by our FEniCS implementation of the above computational model.

#### S6 Supplemental Videos

**Supplemental Video 1 (*Experiment and data analysis*).** This video shows the tracking and reconstruction of a flower of the primary inflorescence in a 29-day-old sample of *Arabidopsis thaliana* (ecotype Col-0) grown under normal gravity conditions (1 g) and continuous light at the SAMBA laboratory of SISSA. The images were acquired by using two digital cameras, namely, two acA4024-29uc c mount cameras from Basler, both of which were equipped with an objective m0824-mpw2 from Computar. The points (black dots) were tracked on the stereo pair of images, which were calibrated to reconstruct the 3D position by exploiting the Computer Vision Toolbox in MATLAB R2019b. The colored lines, from blue to red for increasing time, are obtained by moving averaging over ten detected positions. The red dot indicates the base of the plant.

**Supplemental Video 2 (*A growing shoot*).** Computational results from the nonlinear model (S4.1) for $\ell_0 = 6.8\,\text{cm}$, $\ell_g = 7\,\text{cm}$, $\alpha = 0.2$, $\beta = 0.8$, $\eta = 20$, $\tau_a = 2\,\text{min}$, $\tau_m = \tau_r = \bar{\tau}_m = \bar{\tau}_r = 12\,\text{min}$, $\tau_e = 24\,\text{min}$, $\tau_g = 40\,\text{h}$, $\tau_\ell = 6\,\text{d}$, $r = 0.5\,\text{mm}$, $\rho = 10^3\,\text{Kg}\,\text{m}^{-3}$, $E_0 = 10^7\,\text{Pa}$, $E_1 = 200\,E_0$, and $\nu = 0.5$. We notice that the critical length for the stability of the corresponding reduced model (S4.5) is given by $\ell^\star \approx 7.24\,\text{cm}$ for the chosen model parameters. Here black arrows denote the statoliths local orientations ($\mathbf{h}$), while the other three are the directors ($\mathbf{d}_1$, $\mathbf{d}_2$ and $\mathbf{d}_3$). In the beginning only endogenous oscillations with gravitropic and proprioceptive responses are observed. In the intermediate regime the oscillations due to the intrinsic oscillator and the mechanical flutter are comparable and give rise to trochoid-like patterns. In the end, the flutter instability dominates the oscillatory movements.

**Supplemental Video 3 (*A growing shoot*).** Same as in Supplemental Video 2 but for $\alpha = 0.01$. Differently from Supplemental Video 2, intrinsic cues have little influence and the growing shoot tends to circular patterns passing by elliptic trajectories.

**Supplemental Video 4 (*Limit cycles*).** Computational results from the nonlinear model (S4.1) with equations (S4.1a)-(S4.1b) replaced by $\lambda \equiv 1$, for $\ell = 6.59\,\text{cm}$, $\alpha = 0$, $\beta = 0.8$, $\eta = 20$, $\tau_m = \tau_r = \bar{\tau}_m = \bar{\tau}_r = 12\,\text{min}$, $\tau_a = 2\,\text{min}$, $\tau_g = 20\,\text{h}$, $r = 0.5\,\text{mm}$, $\rho = 10^3\,\text{Kg}\,\text{m}^{-3}$, $E_0 = E_1 = 10^7\,\text{Pa}$, and $\nu = 0.5$. Here black arrows denote the statoliths local orientations ($\mathbf{h}$), while the other three are the directors ($\mathbf{d}_1$, $\mathbf{d}_2$ and $\mathbf{d}_3$). The model is initially perturbed with an apical load $\epsilon \sin(\pi\, t/12)\,(2\mathbf{e}_1 + \mathbf{e}_3)$ where $\epsilon = 10^{-5}\,\text{N}$ for $t \in [0, 12]$ min. Upon such a perturbation, the solution tends to pendular oscillations until an orthogonal apical load $\epsilon \sin(\pi(t - 1800)/12)\,(\mathbf{e}_1 - 2\mathbf{e}_3)$ is applied for $t \in [1800, 1812]$ min. Then the solution tends to the stable circular limit cycle by describing elliptic spirals.

**Supplemental Video 5 (*Epitrochoid*).** Same model as in Supplemental Video 4, but for $\alpha = 0.3$, $\tau_e = 20\,\text{min}$ and $\ell = 6.565\,\text{cm}$. In order to speed up the convergence to periodic oscillations, some perturbations were introduced as apical loads at different time intervals (not shown).

**Supplemental Video 6 (*Hypotrochoid*).** Same as in Supplemental Video 5 but with opposite angular velocity for the internal oscillator.

[17] J. von Sachs. Über orthotrope und plagiotrope Pflanzentheile. *Arbeiten des Botanischen Instituts in Würzburg*, 2:226–284, 1882.
